# Development and Application of Home Cage Monitoring in Laboratory Mice and Rats: a Systematic Review

**DOI:** 10.1101/2023.03.07.531465

**Authors:** Pia Kahnau, Paul Mieske, Jenny Wilzopolski, Otto Kalliokoski, Silvia Mandillo, Sabine M. Hölter, Vootele Voikar, Adriana Amfim, Sylvia Badurek, Aleksandra Bartelik, Angela Caruso, Maša Čater, Elodie Ey, Elisabetta Golini, Anne Jaap, Dragan Hrncic, Anna Kiryk, Benjamin Lang, Natasa Loncarevic-Vasiljkovic, Hamid Meziane, Aurelija Radzevičienė, Marion Rivalan, Maria Luisa Scattoni, Nicolas Torquet, Julijana Trifkovic, Brun Ulfhake, Christa Thöne-Reineke, Kai Diederich, Lars Lewejohann, Katharina Hohlbaum

## Abstract

Traditionally, in biomedical animal research, laboratory rodents are individually examined in test apparatuses outside their home cages at selected time points. However, the outcome of such tests can be influenced by the novel environment, the time of day, separation from the social group, or the presence of an experimenter. Moreover, valuable information may be missed when the animals are only monitored in short periods. These issues can be overcome by longitudinal monitoring mice and rats in their home cages. To shed light on the development of home cage monitoring (HCM) and the current state of the art, a systematic review was carried out on 521 publications retrieved through PubMed and Web of Science. Both the absolute (∼ ×26) and relative (∼ ×7) number of HCM-related publications increased from 1974 to 2020. In both mice and rats, there was a clear bias towards males and individually housed animals, but during the past decade (2011–2020), an increasing number of studies used both sexes and group housing. More than 70 % of the studies did not involve a disease model, but the percentage of studies using disease models increased since the 2000s. In most studies, animals were kept for short (up to 4 weeks) length periods in the HCM systems; intermediate length periods (4–12 weeks) increased in frequency in the years between 2011 and 2020. Before the 2000s, HCM techniques were predominantly applied for less than 12 hours, while 24-hour measurements have been more frequently since the 2000s. The systematic review demonstrated that manual monitoring is decreasing but still relevant. Until (and including) the 1990s, most techniques were applied manually but have been progressively replaced by automation since the 2000s. Independent of the publication year, the main behavioral parameters measured were locomotor activity, feeding, and social behaviors; the main physiological parameters were heart rate and electrocardiography. External appearance-related parameters were rarely examined in the home cages. Due to technological progress and application of artificial intelligence, more refined and detailed behavioral parameters could be investigated in the home cage in recent times.

Over the period covered in this study, techniques for HCM of mice and rats has improved considerably. This development is ongoing and further progress and validation of HCM systems will extend the applications to allow for continuous, longitudinal, non-invasive monitoring of an increasing range of parameters in group-housed small rodents in their home cages.

## Introduction

In biomedical research, laboratory rodents are traditionally removed from their home cages for a defined period of time (from a few minutes up to several hours) and placed in experimental apparatuses to measure parameters of interest. The experimental apparatuses are usually unfamiliar to the animals and, therefore, represent a novel environment. In several cases, experimental designs include familiarization sessions to reduce the potential impact of emotional factors before collecting the parameters of interest. Common examples for monitoring behavioral parameters such as anxiety-related behavior or locomotor activity are the Open Field and the Elevated Plus Maze tests [1]. Both tests are based on mice’ and rats’ natural avoidance of open areas (Open Field) where risk of predation is high and unsafe enclosure (height with no protection in the Elevated Plus Maze), measuring exploration and locomotor activity as expressions of anxiety-like behavior [2]. However, such brief tests, performed outside of the home cage, may not adequately reflect the complexity of, for example, anxiety-like behavior [3–7]. Moreover, the behaviors displayed by an animal in these tests depend also on unrelated factors [8] and it has been shown that the reproducibility of test results obtained in this way is low [9].

The same applies to the collection of other than behavioral data outside the home cage. If, for instance, body temperature is measured using a rectal probe, an animal will generally be removed from its home cage and be hand-restrained for the duration of the measurements, which can result in a stress response and stress-associated hyperthermia [10].

When the animals are tested outside their home cage, the time of day can play a critical role, too. Although rats and mice are nocturnal/crepuscular animals, experiments are often carried out during the daytime when the lights are turned on in the animal and experimental rooms [11–13]. In addition, it needs to be considered which behaviors are observed at what time. Mice show higher activity in an Open Field test during the dark phase when compared to the light phase [14]. Mice also consume less food during the light phase and are less willing to work for access to food in comparison to their willingness during the dark phase [15].

Other disadvantages of tests performed outside the animals’ home cage are that the animals are handled by an experimenter and separated from their familiar social group, which may negatively influence both animal welfare and data quality [16–19]. The lack of reproducibility in animal-based research has been a major issue for a long time. In 1999, it was demonstrated that, in spite of planning and stringent compliance with protocols, different laboratories obtained different results for the same experiments [3]. To date, the reproducibility crisis remains unresolved and various approaches are discussed and pursued to improve the reliability of scientific data [3, 7, 20–24]. One approach that can be used to minimize variability between laboratories is to monitor the animals in their familiar environment (*i.e.*, in their home cage). Housing conditions for laboratory animals vary considerably between animal facilities around the world and there is no standard definition of a home cage. However, a home cage can generally be considered as the environment where the animals spend most of their lifetime [25, 26]. In the European Union this environment must meet the minimum requirements defined in the Directive 2010/63/EU. The Directive stipulates that social animals should be kept in groups whenever possible [27]. The minimum floor area of a home cage should be adjusted depending on the species, the number of animals, and their weight it houses. Food, water, bedding, and nesting material must be provided [27]. Furthermore, the home cage should be structured in a way that allows a wide range of natural behaviors to be exhibited [27]. In the home cage, the animals are usually undisturbed and behave in a way that can be considered “natural” under laboratory husbandry conditions.

A broad range of parameters can be monitored in the home cage using manual or automated techniques, *e.g.*, a variety of behavioral parameters such as activity [28, 29], social behavior [30–32], learning and memory [33–35], feeding [36–38] and physiological parameters like heart rate or body temperature [39–42]. Manual techniques used for home cage monitoring (HCM) include live observations or analyses of videos recorded by camera-based systems, which can be complemented or substituted by automated HCM techniques. Automated HCM techniques have been implemented in various commercial systems. A range of systems were described in a review by Richardson *et al.* [43]. For example, a combination of light emitting diodes (LEDs), infrared light sensors, and a camera allows for analyzing, among others, physical activity or learning behavior [44, 45]. Electrical capacitance technology makes it possible to measure physical activity or rest [46, 47]. As animals move about in a cage, changes in an emitted weak electromagnetic field can be used to track their movements. However, these systems do not allow the acquisition of individual data from group-housed animals. Data from individuals, also when they are kept in social groups, can be generated using radio-frequency identification (RFID) systems. For instance, the activity of 20–40 mice can be automatically measured in a large semi-naturalistic home cage [48, 49]. RFID systems enable to test learning and memory of mice and rats using operant conditioning corners that grant or deny access to water [35, 50, 51] and to perform preference tests in the home cage [52, 53]. RFID systems can be combined with other techniques such as depth-sensing infrared camera for automatic individual tracking of animals [54].

HCM offers several advantages for both data quality [55–57] and animal welfare [46, 58, 59]. Ideally, group-housed animals are observed and/or tested in their familiar environment 24/7 and remain undisturbed by the experimenters. HCM has grown in importance and demands have changed over the last decades driven by technology and digitalization. However, automated 24-hour measurements for long-term periods still seem to be a major challenge. In 2021, European researchers joined forces in a project titled “Improving biomedical research by automated behaviour monitoring in the animal home-cage” (TEATIME: cost-teatime.org) in order to promote the further use and development of HCM systems. On this website, a comprehensive catalog of currently existing HCM systems is provided (cost-teatime.org/about/technologies). Approximately at the same time, another initiative called “Translational Digital Biomarkers” was launched within the North American 3Rs Collaborative (www.na3rsc.org/tdb). To identify past trends in and the current state of HCM, a systematic review was carried out in close collaboration with the COST Action TEATIME network. The following research questions were addressed: How have techniques and applications for home cage monitoring of laboratory mice and rats developed over time? Has the degree of automatization for monitoring behavioral, physiological, and external appearance-related parameters changed?

## Materials & Methods

### Review protocol

The protocol of the systematic review was uploaded to the Open Science Framework on May 3, 2021 after the search for literature was conducted: https://osf.io/4gzcx. It was registered on March 14, 2022 when the full text review and extraction phase were in progress: https://osf.io/un5e6.

### Search strategy and screening

Primary databases were searched through PubMed and Web of Science on February 24, 2021. PubMed search terms were refined using the search refiner tool QueryVis [60]. The following search strings were used for obtaining relevant studies.

Pubmed: ((“mice”[MeSH Terms] OR mice OR mouse) OR (“rats”[MeSH Terms] OR rat OR rats)) AND ((home cage) OR (housing cage)) AND (monitoring OR (“observation”[MeSH Terms] OR observation)). The filter “other animals” was applied.

Web of Science: ALL FIELDS: (((mice OR mouse) OR (rats OR rat)) AND ((“home cage”) OR (“housing cage”)) AND (monitoring OR observation)).

The data extraction template can be found in S1 Supporting Information. Further detailed information on the objective and materials & methods is provided in S2 Supporting Information. In brief, only primary studies (published in English language) in which behavioral, physiological, and/or external appearance-related parameters were monitored in mice and/or rats in their home cage or a testing apparatus connected to the home cage were included. Studies in which no other parameters than calorimetric measurements were examined in a home cage were excluded. In phase 1, titles and abstracts were screened. Thereafter, full texts were screened (phase 2) and data were extracted (phase 3) simultaneously. In all phases, all papers were screened by two independent reviewers and discrepancies were resolved by a third reviewer.

### Analysis

The responses for the mouse/rat strain were reviewed by two persons: In mice, the 129 substrains were subsumed under “129”, C57BL/6J and C57BL/6N under “C57BL/6”, and ICR and CD-1 under “CD-1”. In rats, Holtzman and Sprague Dawley were subsumed under “Sprague Dawley”. For genetically modified animals, the background strain was extracted. If strains were on a mixed genetic background, “mixed background” was indicated.

If responses appeared not to be plausible or free text-responses had to be clustered, the answers were corrected and the corrected responses were used for further analysis. All original and corrected responses can be found in S6 Data.

SPSS (IBM Corp. Released 2020. IBM SPSS Statistics for Windows, Version 27.0. Armonk, NY: IBM Corp) was used for the linear regression analysis and for creating the figures. In the figures visualizing the historical change, the year 2021 was excluded since the literature search was carried out in February 2021 and did not cover the entire year. The ratio of relevant HCM references per year was analyzed using a linear regression model. The R^2^ (fraction of explained variance) indicates how well the model fits the data (from 1, best prediction to 0, no prediction). The remaining data were descriptively analyzed.

## Results

### Overview of the studies

The searches through Pubmed and Web of Science retrieved 1079 and 341 references, respectively. Figure 1 shows the PRISMA flow diagram. A total of 241 duplicates were removed. Titles and abstracts of 1179 references were screened, of which the full text for 721 references were assessed for eligibility. 521 references were retained for data extraction.

**Figure 1.**
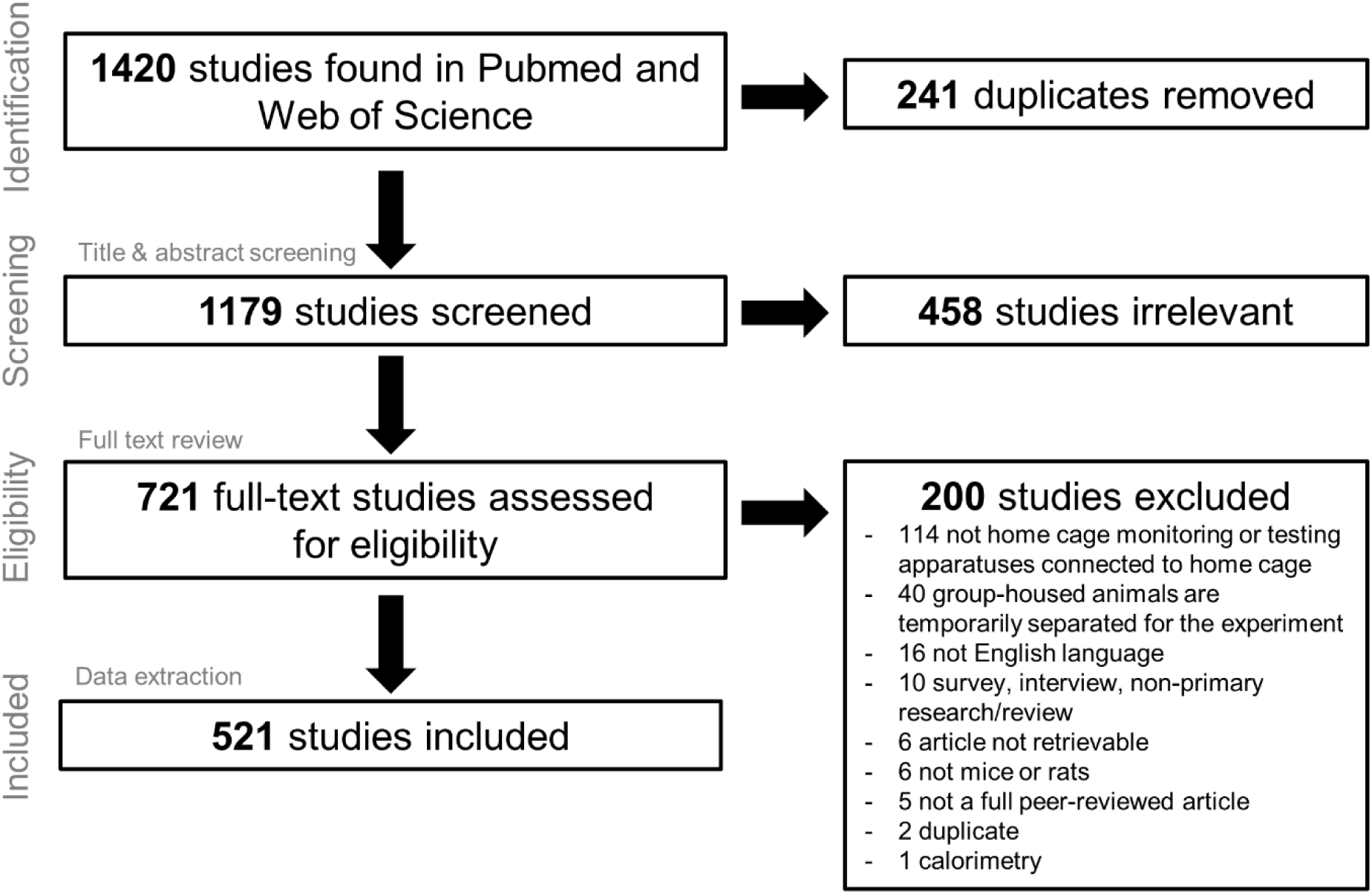
PRISMA flow diagram.

### Number of publications on HCM

The earliest publication fulfilling our criteria for home cage monitoring, that we could find, was published in 1974. The yearly number of publications in which mice and/or rats were monitored in their home cages according to our eligibility criteria increased (Figure 2A). In 1987, no publication meeting the inclusion criteria was found.

**Figure 2.**
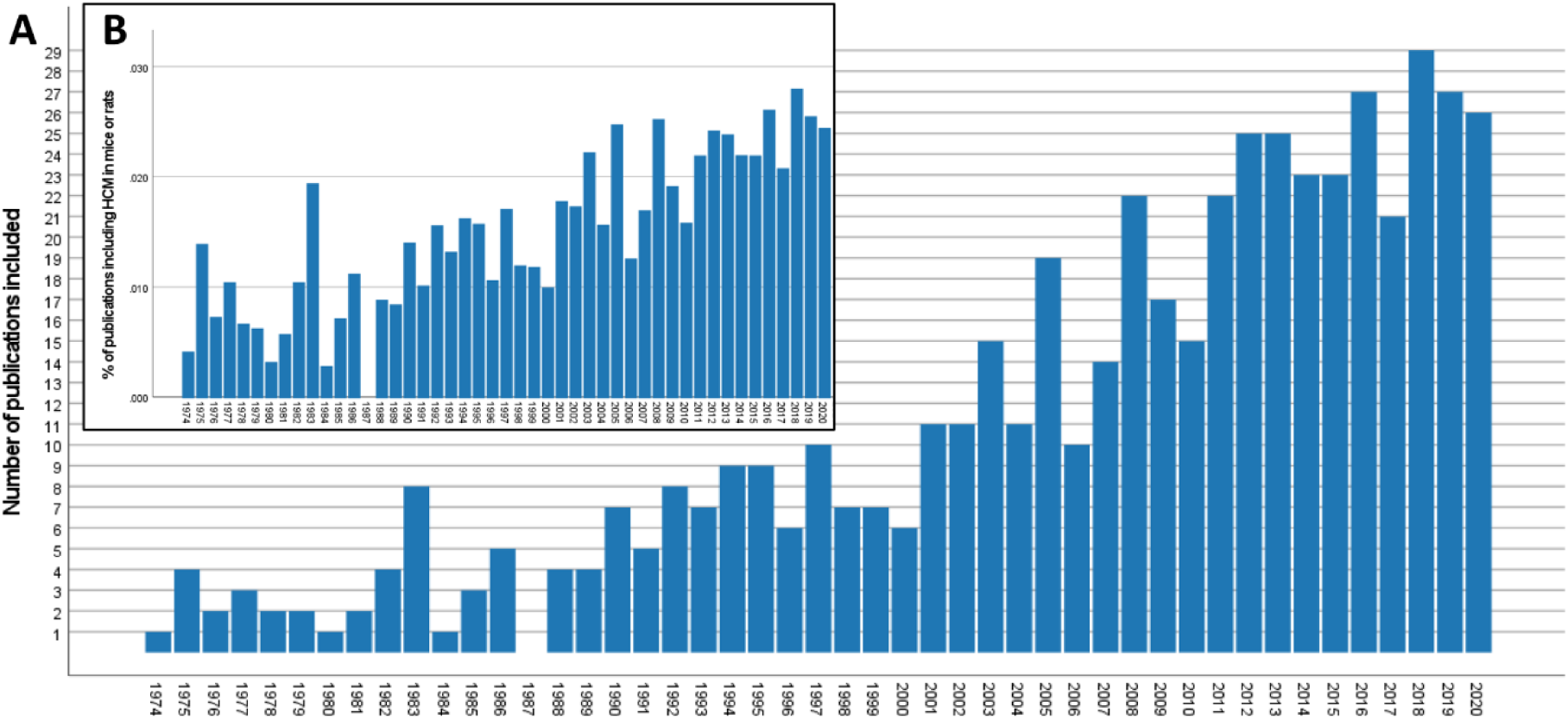
Historical change in the publication rate. (A) Absolute number of included publications on home cage monitoring (HCM) of mice and rats. (B) The ratio of HCM references relative to the overall number of studies in mice or rats listed in PubMed published in the years 1974 to 2020.

When comparing the number of HCM publications with the overall number of studies in mice or rats listed in PubMed (search string: “mice”[MeSH Terms] OR “mice”[All Fields] OR “rats”[MeSH Terms] OR “rats”[All Fields]; filter: other animals), the percentage of relevant HCM references increased over time (Figure 2B; linear regression analysis: F(1, 45) = 100.704, p < 0.001; R^2^ = 0.691).

The first study was dated in 1974; in 2019 and 2020, 29 and 26 publications were identified, respectively. In 2021, only one study fulfilled the eligibility criteria, but since the literature search was carried out in February 2021 the databases did not cover the entire year.

### Species and sex

Mice were studied in 276 publications, rats in 240, and both species in 5 publications (Table 1). Most studies included males. The percentage of studies using exclusively males was 55 % in mice (13 % females only), 69 % in rats (11 % females only) and 40 % in studies of both species (20 % females only). For the past 10 years (2011– 2020), the percentage of included studies using animals of both sexes increased (Figure 3A).

**Figure 3.**
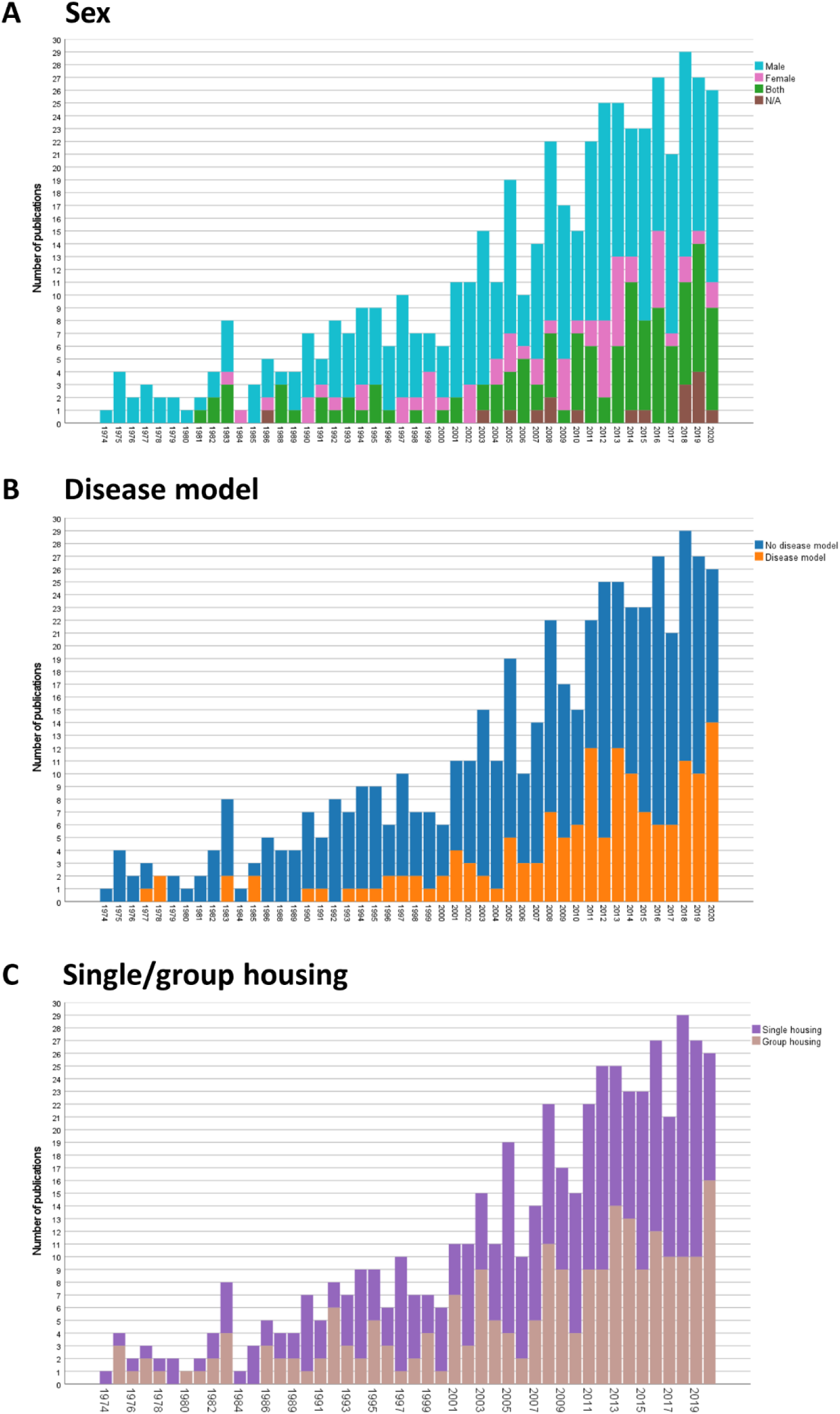
Historical change in the number of publications using male and/or female mice (A), publications investigating specific disease models (B), and publications employing single or group housing (C).

**Table 1.**
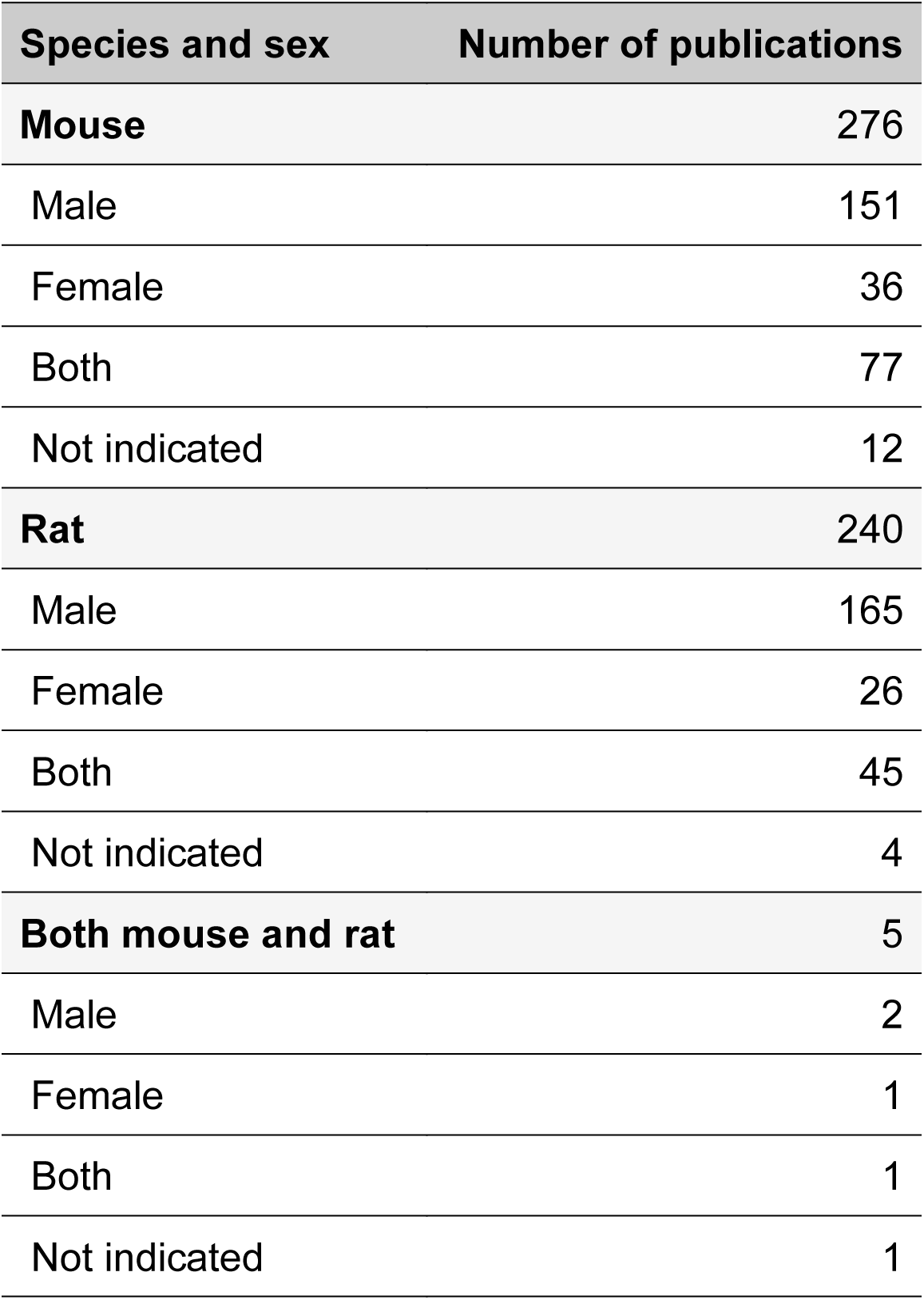
Species and sex of animals used in the included publications.

| Species and sex | Number of publications |
| --- | --- |
| **Mouse** | 276 |
| Male | 151 |
| Female | 36 |
| Both | 77 |
| Not indicated | 12 |
| **Rat** | 240 |
| Male | 165 |
| Female | 26 |
| Both | 45 |
| Not indicated | 4 |
| **Both mouse and rat** | 5 |
| Male | 2 |
| Female | 1 |
| Both | 1 |
| Not indicated | 1 |

### Strains

The most frequently used mouse strains were C57BL/6 (n = 192), BALB/c (n = 31), CD-1 (n = 26), DBA/2 (n = 21), and 129 (n = 16), whereas the most frequently used rat stocks were Sprague Dawley (n = 115), Wistar (n = 74), and Long Evans (n = 34) (Figure 4). Among mouse strains inbred are most common while among rats the reverse is true. In S3 Supporting Information, the numbers of publications including particular strains, even those subsumed under “other” in Figure 4, are listed.

**Figure 4.**
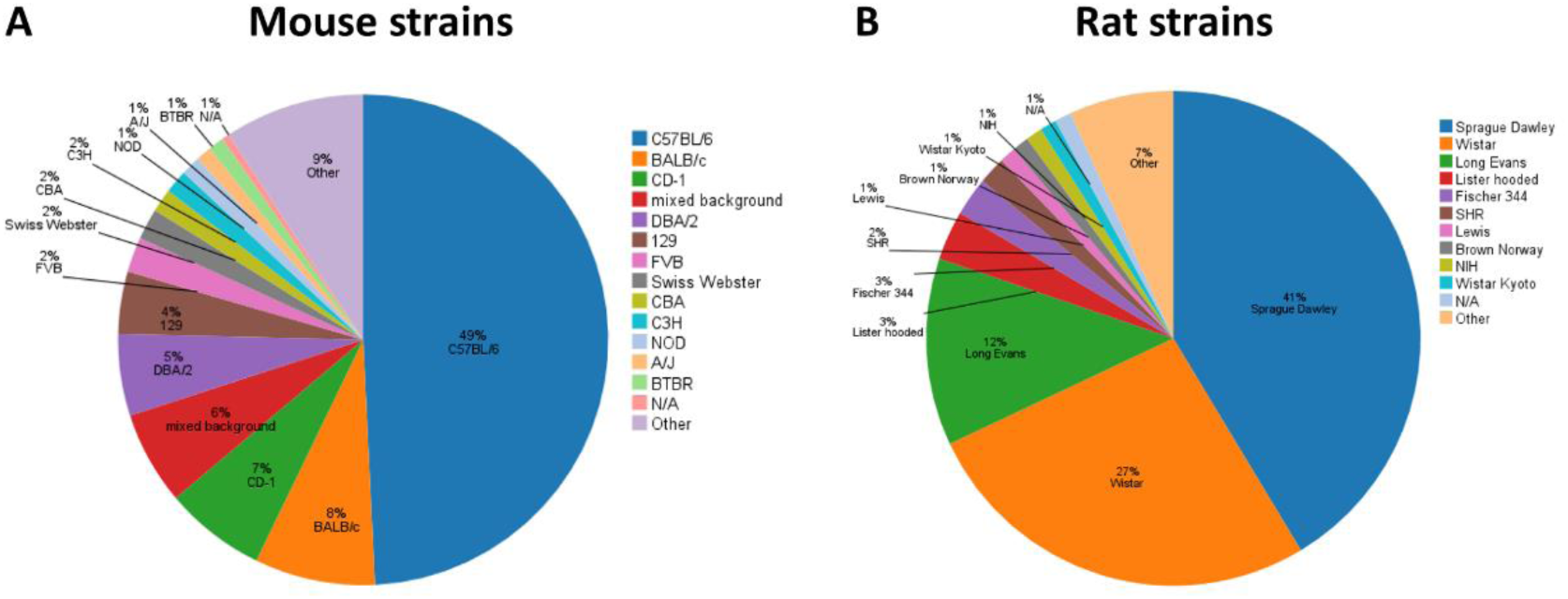
Mouse and rat strains that were used in the included publications in the years 1974 to 2021. More than one strain could be used in a study. Strains that were indicated twice or once only, were summarized under “other”.

### Disease models

The disease models were categorized according to ICD-11 (https://icd.who.int/browse11/l-m/en). The most frequently studied disease models in the included references were mental, behavioral or neurodevelopmental disorders (n = 74), diseases of the nervous system (n = 32), and endocrine, nutritional or metabolic diseases (n = 15; Table 2). In a majority of publications – 367 references – no disease model was studied. However, the percentage of included studies involving disease models increased in the two past decades (Figure 3B; 1974–1980: 20 %; 1981–1990: 13 %; 1991–2000: 18 %; 2001–2010: 27 %; 2011–2020: 38 %).

**Table 2.** Disease models used in the included publications. In one paper, more than one disease model was studied. The disease models were categorized according to ICD-11 (https://icd.who.int/browse11/l-m/en).

| Disease model | Number of publications |
| --- | --- |
| No disease model | 368 |
| Mental, behavioral or neurodevelopmental disorders | 74 |
| Diseases of the nervous system | 32 |
| Endocrine, nutritional or metabolic diseases | 15 |
| Diseases of the circulatory system | 8 |
| Diseases of the digestive system | 5 |
| Diseases of the immune system | 4 |
| Infectious or parasitic diseases | 4 |
| Developmental anomalies | 2 |
| Diseases of the respiratory system | 2 |
| Injury, poisoning or certain other consequences of external causes | 2 |
| Neoplasms | 2 |
| Sleep-wake disorders | 2 |
| Symptoms, signs or clinical findings, not elsewhere classified | 1 |
| Diseases of the musculoskeletal system or connective tissue | 1 |

### Home cage monitoring systems

S4 Supporting Information provides detailed information about the HCM systems used in the included studies. The table contains the number of publications in which the HCM systems were used, and the species, strain/stock, sex, and group size of the animals housed in the systems. Moreover, the duration the animals spent in the particular system is listed. In 126 cases, custom-built HCM systems and in 217 cases commercially available HCM systems were used (multiple HCM systems were applied in 28 studies). A “custom-built” system had to fulfill two criteria: 1) it is not commercially available and construction plans and/or software was provided by the authors; 2) the authors gave a name to the system. If these criteria were not met and neither a commercially available nor a “custom-built” system was used, it was considered that no system was applied in the study, *e.g.*, the deployment of a camera only. The use of commercially available HCM systems has increased over time (Figure 5).

**Figure 5.**
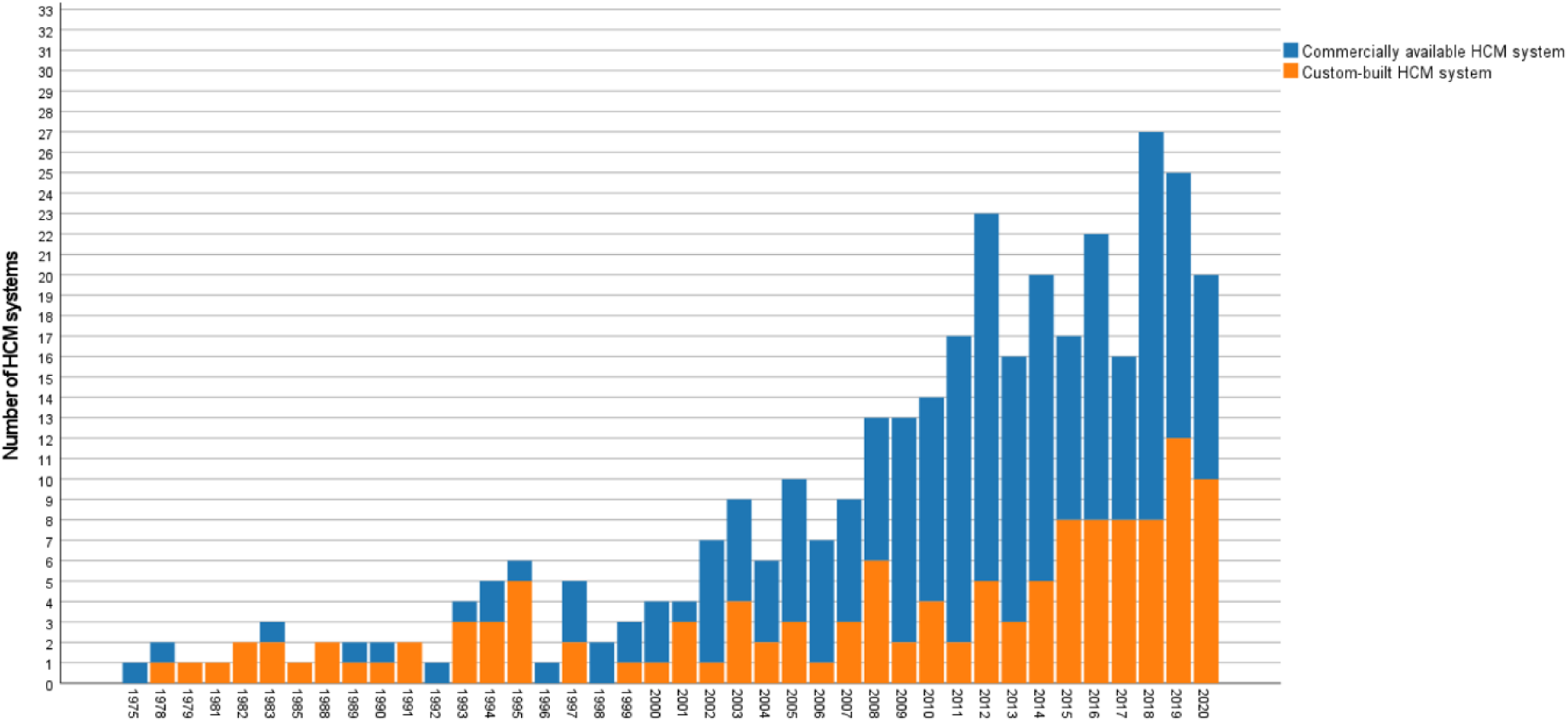
Historical change in the number of custom-built and commercially available home cage monitoring (HCM) systems. More than one system could be applied in a study.

### Social housing conditions

In most studies, animals were individually housed in their home cage system (n = 298) (Table 3). In the 1970s, there were as many studies in which animals were kept individually as in groups. After 1980, group housing has decreased in comparison to the 1970s, but there was a slight increasing trend between the 1980s and the 2010s (Figure 3C; 1974–1980: 47 %, 1981–1990: 32 %, 1991–2000: 31 %, 2001–2010: 33 %, 2011–2020: 38 %). In contrast, the number of studies with single-housed animals has increased after 1980 in comparison to the 1970s and slightly decreased again in the 2000s and 2010s (1974–1980: 47 %, 1981–1990: 61 %, 1991–2000: 61 %, 2001–2010: 59 %, 2011–2020: 54 %). Table 3 shows that those laboratory rodents that were socially housed were often kept in pairs (n = 60) more so than in groups of three (n = 27) or four animals (n = 30). Other group sizes between five and nine animals were even less frequently found. Groups of ten (n = 12) or even more than ten animals (n = 16) were rarely reported in the included publications.

**Table 3.** Number of animals housed together in a cage.

| Number of animals housed in a home cage system | Number of publications |
| --- | --- |
| 1 | 298 |
| 2 | 60 |
| 3 | 27 |
| 4 | 30 |
| 5 | 15 |
| 6 | 9 |
| 7 | 2 |
| 8 | 8 |
| 9 | 6 |
| 10 | 12 |
| More than 10 | 16 |
| Not indicated | 38 |

Considering housing conditions for single-sex studies, 68 % of male mice were individually housed, while only 36 % of the female mice were kept in social isolation (Table 4). In contrast to male mice, most female mice (61 %) were kept in groups. For rats, both sexes (62 % of the males and 65 % of the females) were predominantly single-housed.

**Table 4.** Percentage of publications in which males and females were housed individually or in groups. For this table, publications in which both species or both sexes of a species were used or the sex of the animals was not indicated were neglected. Due to rounding, the numbers do not always sum to 100 %

|  | Individual housing<br>(%, abs. number<br>given in brackets) | Group housing<br>(%, abs. number<br>given in brackets) | Not indicated<br>(%, abs. number<br>given in brackets) |
| --- | --- | --- | --- |
| **Mice** |  |  |  |
| Males | 68 (102) | 26 (39) | 7 (10) |
| Females | 36 (13) | 61 (22) | 3 (1) |
| **Rats** |  |  |  |
| Males | 62 (103) | 33 (55) | 4 (7) |
| Females | 65 (17) | 23 (6) | 12 (3) |

### Duration of housing in the home cage

The duration of housing was classified as short-term (1–28 days), intermediate length (1–3 months), or long-term (more than 3 months). Regardless of the publication year, researchers seemed to favor short-term housing (2–7 days: n = 75; 1–2 weeks: n = 73; 2–4 weeks: n = 102). Intermediate duration housing was not uncommon (4–12 weeks; n = 127; Figure 6A). However, single-day housing for HCM (n = 11) and longer periods (12–24 weeks, n = 35; 24–48 weeks, n = 8; more than 48 weeks, n = 7) were rare. In 16 % of the studies (n = 83), the authors did not indicate the duration of housing.

**Figure 6.**
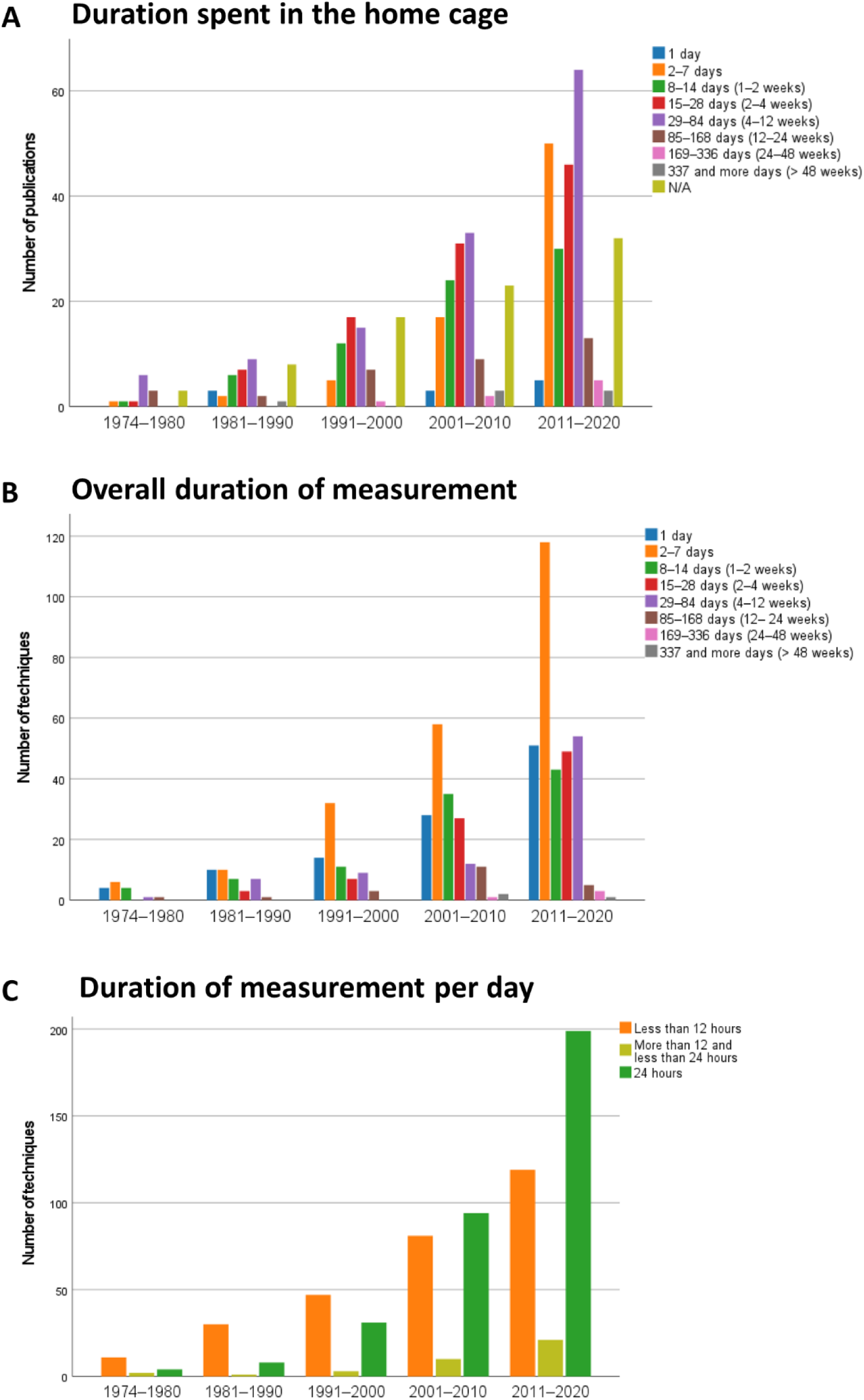
Historical change in the duration of housing and monitoring in the home cage: (A) duration spent in the home cage, (B) overall duration of measurement, (C) duration of measurement per day (C) using a home cage monitoring technique. For (B) and (C), it must be noted that more than one technique for data measurement could be used in a study (*i.e.*, multiple answers per study for each technique are possible). If one of the techniques used to investigate parameters of interests could not be extracted from the paper, this technique was excluded for (B) and (C). Note the differences in the scaling of the y-axis.

### Overall duration of monitoring in the home cage

Figure 6B gives information on how long a technique was used for measuring behavioral, physiological, or external appearance–related parameters. In all years, most studies involved measurements of 2–7 years. In 1974–1980, the animals were mostly monitored for 1–14 days. Only in rare cases did the measurements last for 4– 24 weeks. In 1981–1990, the proportion of studies in which parameters were analyzed for 2–12 weeks increased. Between 2011 and 2020, the frequencies of measurements over 1 day, 1–2 weeks, 2–4 weeks, and 4–12 weeks were relatively similar, with the latter being clearly increased in comparison to the two past decades. Longer periods of measurement (*i.e.*, > 3 months) were only found in a few cases.

There was a slight change over time in the relation between the housing duration in the HCM systems and the duration for which the parameters of interest were measured. It has become more common to do measurements during the entire time in which the animals are kept in a HCM system over time (1974–1980: 29 %; 1981–1990: 44 %; 1991–2000: 42 %; 2001–2010: 47 %; 2011–2020: 54 %).

### Duration of monitoring per day in the home cage

As shown in Figure 6C, in the time period 1974–2000, most measurements lasted for less than 12 hours per day. Past the year 2000, 24-hour measurements were conducted more frequently.

Overall, automated techniques dominated for measurements of more than 12 hours. For monitoring intervals of less than 12 hours manual techniques were common (70 % of studies).

### Home cage monitoring techniques

In most studies, one HCM technique was used (n = 362). The second most frequently reported options were two techniques (n = 118). A combination of three (n = 35) or more (n = 6) techniques was less often applied.

Table 5 shows that in a group housing setting more techniques were applied to measure data from individuals than from groups of animals. The degree of automatization played a negligible role for the measurement of data from individual subjects, but data from groups of animals were rather generated by manual techniques.

**Table 5.** Percentage of automatic and manual techniques measuring data from individuals and groups of animals in a group-housed setting. Due to rounding, the sum across the cells does not equal 100 %. More than one technique could be applied in a study and the same techniques could be used across the different studies.

|  | **Automatic**<br>(%, abs. number<br>given in brackets) | **Manual**<br>(%, abs. number<br>given in brackets) |
| --- | --- | --- |
| Data from individuals | 33 (83) | 36 (91) |
| Data from groups of<br>animals | 10 (26) | 18 (45) |
| Both data from<br>individuals and<br>groups of animals | 0 (1) | 0 (0) |
| Not indicated | 0 (1) | 1 (3) |

Figure 7A illustrates how the use of techniques for home cage monitoring of laboratory mice and rats has developed over time. Across the whole study periods, manual evaluation (including live monitoring and manual evaluation of videos from RGB or infrared cameras) was the most frequently used technique. In the 2010s, non-invasive methods (*i.e.,* beam-based tracking and visual object tracking) have overtaken the use of telemetry, which are extremely invasive.

**Figure 7.**
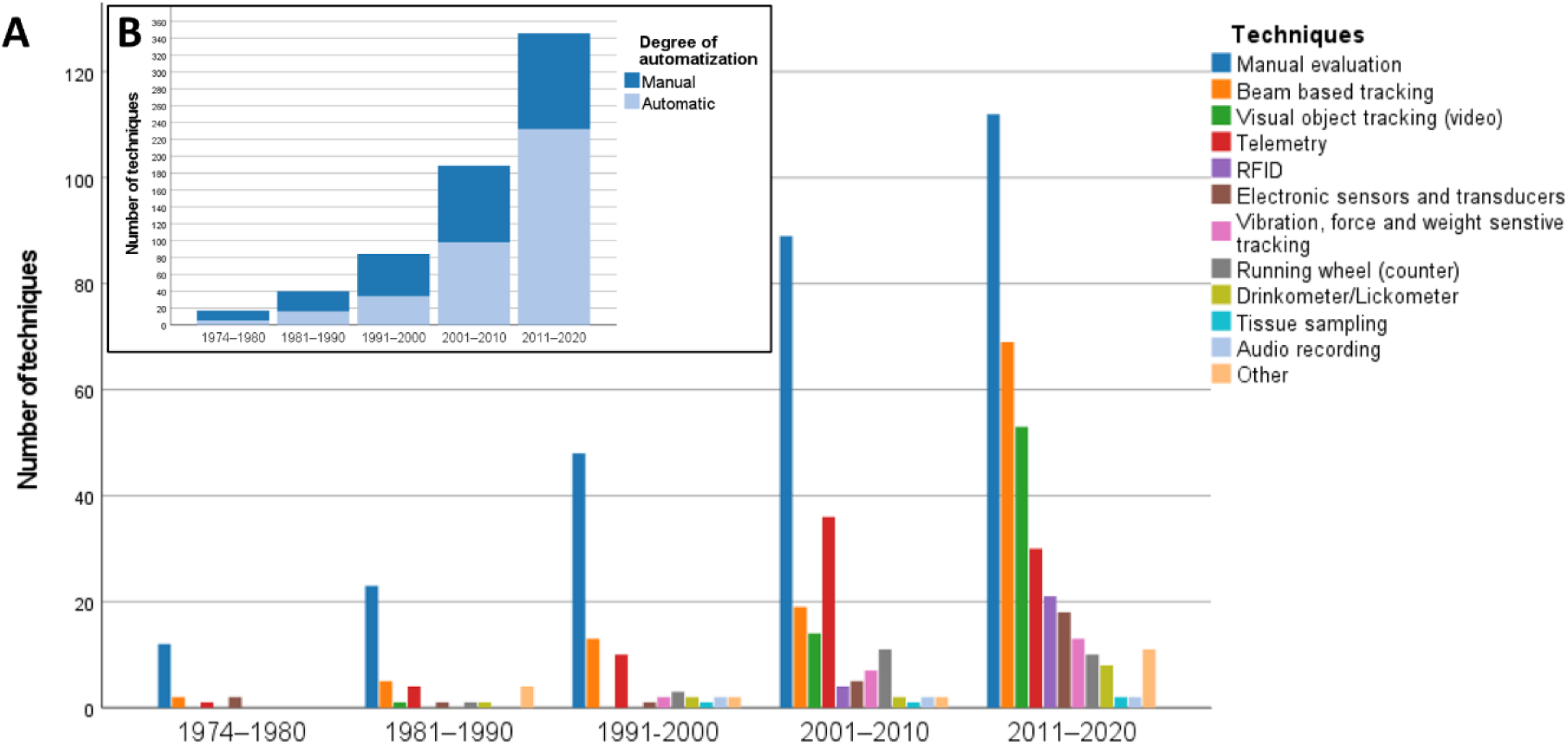
Historical change in home cage monitoring techniques for laboratory mice and rats. More than one technique could be indicated in a publication. (A) Number of techniques used in the included publications in the defined time periods. Other: infrared thermometer (n = 1), automatic food dispenser (n = 1), impedance pneumography (n = 1), flowmeter circuit (n = 1), lever (n = 2), brain imaging cameras (n = 2), fiber photometry system (n = 1), cardiotachometer (n = 1), thermal imaging (n = 1). (B) Number of manual and automated techniques that were applied in the included publications.

The techniques listed in Table 6 were used in group-housed animals and almost all of them allowed for generating data for individual animals. However, it must be considered that often two or more techniques were combined, which may allow for collecting data from individuals instead of animal groups only (*e.g.*, by combining a RFID system, which identifies individual animals, with other techniques).

**Table 6.**
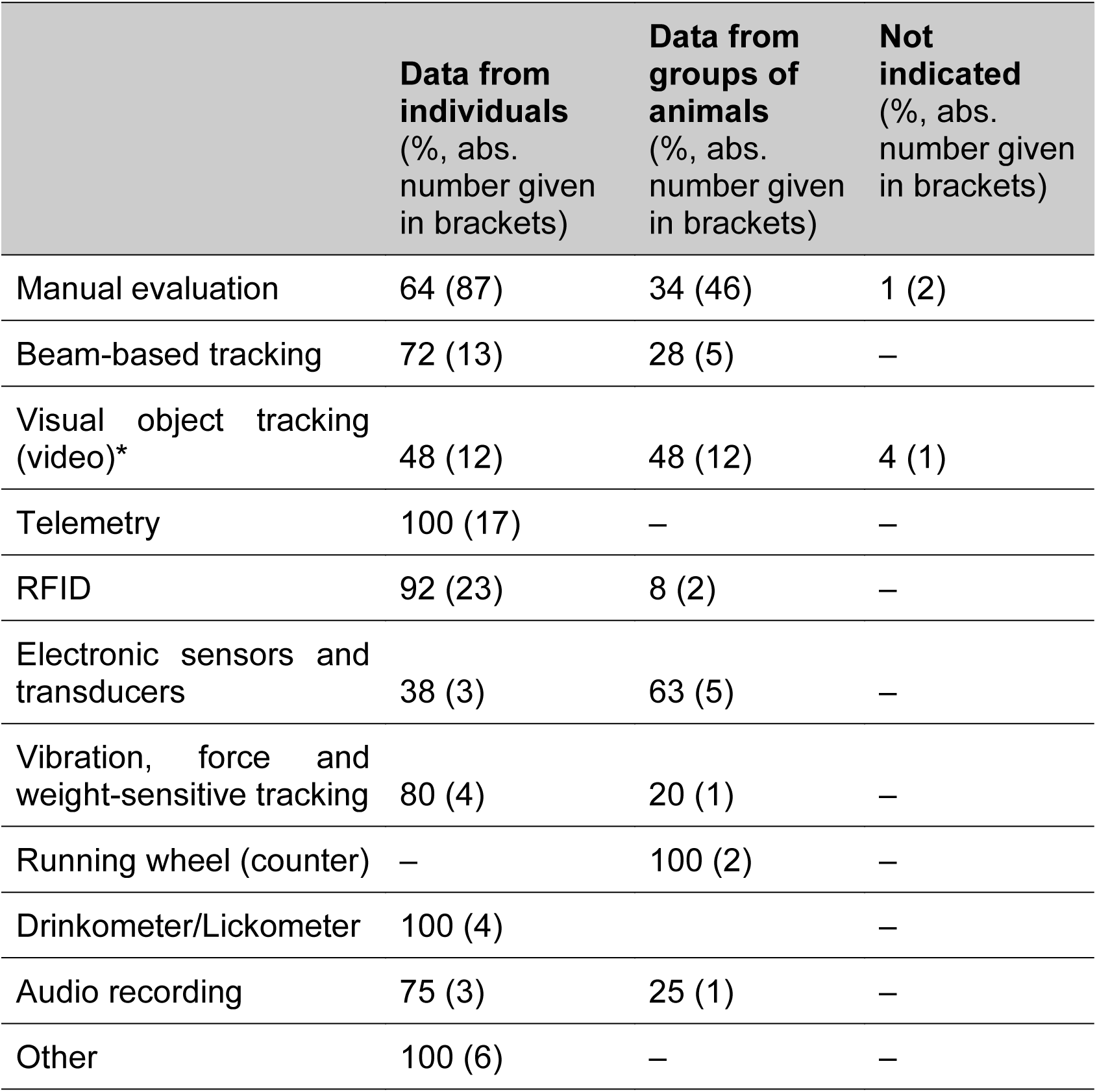
Percentage of techniques that were applied for measuring data from individuals or groups of animals in a group-housed setting. More than one technique could be applied in a study. Due to rounding, the sum across a row does not always equal 100 %. *In one study, this technique was applied for measuring both data from individuals and groups of animals.

|  | **Data from individuals**<br>(%, abs. number given in brackets) | **Data from groups of animals**<br>(%, abs. number given in brackets) | **Not indicated**<br>(%, abs. number given in brackets) |
| --- | --- | --- | --- |
| Manual evaluation | 64 (87) | 34 (46) | 1 (2) |
| Beam-based tracking | 72 (13) | 28 (5) | – |
| Visual object tracking (video)* | 48 (12) | 48 (12) | 4 (1) |
| Telemetry | 100 (17) | – | – |
| RFID | 92 (23) | 8 (2) | – |
| Electronic sensors and transducers | 38 (3) | 63 (5) | – |
| Vibration, force and weight-sensitive tracking | 80 (4) | 20 (1) | – |
| Running wheel (counter) | – | 100 (2) | – |
| Drinkometer/Lickometer | 100 (4) |  | – |
| Audio recording | 75 (3) | 25 (1) | – |
| Other | 100 (6) | – | – |

In the periods between 1974–1980, 1981–1990, and 1991–2000, most measurement were performed manually (71 %; 57 %; 59 %) (Figure 7B). In contrast, between 2001– 2010 and 2011–2020 (52 % and 68 %), most measurements were automated.

In tables 7, 8, and 9, the techniques are separated into methods for assessing behavior, physiology, and external appearance. Manual evaluation and visual object tracking (video) played a role for almost all behavioral parameters (Table 7). Among other techniques, beam-based tracking and RFID were used to analyze locomotor activity, wheel running, motor and sensory functions, feeding, social behavior, as well as learning and memory. RFID systems were used for studying spatial preference, and abnormal behavior. Less frequently used techniques can be found in Table 7.

**Table 7.** Techniques used for the analysis of behavioral parameters. More than one parameter and/or technique could be indicated in a publication. Note that in some cases RFID systems are combined with other techniques to identify the animals before a parameter is measured.

| **Behavioral parameters** | **Number of indications over all studies** |
| --- | --- |
| **Locomotor activity** (*in 1 study, 2 techniques were used) | **317** |
| Manual evaluation | 91 |
| Beam-based tracking* | 81 |
| Visual object tracking (video)* | 54 |
| Telemetry | 48 |
| Electronic sensors and transducers | 13 |
| Vibration, force and weight-sensitive tracking | 12 |
| RFID (radio-frequency identification) | 11 |
| Not indicated | 7 |
| **Feeding (drinking, food)** (*in 2 studies, 2 techniques were used) | **213** |
| Manual evaluation | 123 |
| Beam-based tracking* | 21 |
| Visual object tracking (video) | 13 |
| Drinkometer/lickometer* | 11 |
| RFID* | 11 |
| Vibration, force and weight-sensitive tracking* | 9 |
| Electronic sensors and transducers | 4 |
| Other (automatic food dispenser) | 1 |
| Not indicated | 20 |
| **Social behavior** | **117** |
| Manual evaluation | 101 |
| Visual object tracking (video) | 11 |
| RFID | 2 |
| Audio recording | 1 |
| Beam-based tracking | 1 |
| Not indicated | 1 |
| **Burrowing and nesting** | **56** |
| Manual evaluation | 47 |
| Visual object tracking (video) | 3 |
| Beam-based tracking | 4 |
| Vibration, force and weight-sensitive tracking | 1 |
| Not indicated | 1 |
| **Wheel running** | **51** |
| Running wheel (counter) | 25 |
| Electronic sensors and transducers | 8 |
| Beam-based tracking | 4 |
| Manual evaluation | 2 |
| RFID | 1 |
| Visual object tracking (video) | 1 |
| Not indicated | 10 |
| **Abnormal behaviors** | **34** |
| Manual evaluation | 31 |
| Electronic sensors and transducers | 1 |
| RFID | 1 |
| Visual object tracking (video), | 1 |
| **Facial expression and body posture** | **29** |
| Manual evaluation | 25 |
| Visual object tracking (video) | 4 |
| **Grooming** | **27** |
| Manual evaluation | 20 |
| Visual object tracking (video) | 4 |
| Vibration, force and weight-sensitive tracking | 3 |
| **Learning and memory** | **25** |
| Manual evaluation | 6 |
| Beam-based tracking | 5 |
| RFID | 4 |
| Drinkometer/lickometer | 2 |
| Electronic sensors and transducers | 1 |
| Visual object tracking (video) | 2 |
| Other (lever) | 2 |
| Not indicated | 3 |
| **Anxiety, depression, and schizophrenia** | **15** |
| Manual evaluation | 10 |
| Visual object tracking (video) | 4 |
| Not indicated | 1 |
| **Sleep behavior** | **12** |
| Visual object tracking (video) | 7 |
| Manual evaluation | 3 |
| Telemetry | 1 |
| Vibration, force, and weight-sensitive tracking | 1 |
| **Vocalization** | **11** |
| Manual evaluation | 6 |
| Audio recording | 5 |
| **Motor and sensory functions** | **9** |
| Visual object tracking (video) | 3 |
| Manual evaluation | 2 |
| RFID | 2 |
| Beam-based tracking | 1 |
| Electronic sensors and transducers | 1 |
| **Spatial preference** | **6** |
| Vibration, force, and weight-sensitive tracking | 2 |
| Manual evaluation | 2 |
| RFID | 1 |
| Visual object tracking (video) | 1 |
| **Defecation and urination** | **5** |
| Manual evaluation | 4 |
| Not indicated | 1 |
| **Sniffing** | **4** |
| Manual evaluation | 3 |
| Visual object tracking (video) | 1 |
| **Seizures** | **3** |
| Manual evaluation | 2 |
| Visual object tracking (video) | 1 |
| **Other (each n = 1)** |  |
| Clinical signs not further specified – manual evaluation;<br>colonic contractility – telemetry;<br>curiosity/alertness – manual evaluation;<br>sneezing – manual evaluation;<br>sign of pain or distress not further specified – manual evaluation;<br>twitches – visual object tracking (video);<br>other: behavior not further specified – visual object tracking (video); manual evaluation |  |
| Legend: More than one parameter and/or technique could be indicated in a publication. Note that in some cases RFID systems are combined with other techniques to identify the animals before a parameter is measured. |  |

**Table 8.** Techniques used for the analysis of physiological parameter. More than one parameter and/or technique could be indicated in a publication. Note that in some cases RFID systems are combined with other techniques to identify the animals before a parameter is measured.

| **Physiological parameter** | **Number of indications over all studies** |
| --- | --- |
| **Heart rate & Electrocardiography** | **49** |
| Telemetry | 44 |
| Electronic sensors and transducers | 1 |
| Manual evaluation | 1 |
| Other (cardiotachometer, non-invasive electrodes) | 3 |
| **Body temperature** | **37** |
| Telemetry | 31 |
| RFID | 3 |
| Manual evaluation | 1 |
| Other (infrared thermometer, thermal imaging) | 2 |
| **Body weight** | **2** |
| Vibration, force, and weight-sensitive tracking | 2 |
| **Blood pressure** | **26** |
| Telemetry | 25 |
| Other (flowmeter circuit) | 1 |
| **Neuronal activity** | **21** |
| Telemetry | 9 |
| Electronic sensors and transducers | 1 |
| Tissue sampling | 1 |
| Other (brain imaging cameras, fiber photometry system, implanted electrodes) | 9 |
| Not indicated | 1 |
| **Respiration** | **8** |
| Manual evaluation | 4 |
| Electronic sensors and transducers | 1 |
| Telemetry | 1 |
| Visual object tracking (video) | 1 |
| Other (impedance pneumography) | 1 |
| **Electromyography** | **7** |
| Telemetry | 4 |
| Electronic sensors and transducers | 1 |
| Other (implanted electrodes) | 1 |
| Not indicated | 1 |
| **(Stress) hormones** | **6** |
| Manual evaluation (from fecal samples) | 3 |
| Tissue sampling | 3 |

Telemetry has been used to record most physiological parameters, except for body weight and hormone levels (Table 8). It must be noted that hormone concentrations were measured using microdialysis, automated blood sampling or in fecal samples. The latter were manually collected from the home cages and analyzed later. In the context of the present systematic review, the collection of fecal samples for subsequent analysis was considered a form of manual evaluation. For the analysis of neuronal activity, telemetry was used in most cases. Respiration were mainly manually assessed. Other techniques such as RFID, electronic sensors and transducers, vibration, force, and weight-sensitive tracking, as well as tissue sampling (*e.g.*, microdialysis) were less frequently used.

External appearance was usually evaluated manually (Table 9).

**Table 9.** Techniques used for the analysis of external appearance-related parameters. More than one parameter and/or technique could be indicated in a publication.

| **External appearance-related parameters** | **Number of indications over all studies** |
| --- | --- |
| **Fur condition** | **6** |
| Manual evaluation | 5 |
| Not indicated | 1 |
| **Wounds** | **6** |
| Manual (live) monitoring | 6 |
| **Body Condition Score** | **4** |
| Manual (live) monitoring | 4 |
| **Piloerection** | **4** |
| Manual (live) monitoring | 4 |
| **Chromodacryorrhea** | **1** |
| Manual (live) monitoring | 1 |
| **External appearance-related parameters not further specified** | **3** |
| Manual (live) monitoring | 1 |
| Visual object tracking (video) | 2 |

Detailed information on the methodology of each behavioral, physiological, and external appearance-related parameter (*i.e.*, the degree of automatization, group *versus* individual monitoring, and the duration of measurement) can be found in S5 Supporting Information.

### Behavioral, physiological, and external appearance-related parameters

Figure 8 illustrates the distribution of behavioral, physiological, and external appearance-related parameters between 1974 and 2020. Categories of behavioral parameters (Figure 8A; a range of behaviors were subsumed under a category) that have been investigated throughout were locomotor activity, feeding, social behavior, abnormal behavior, facial expressions and body posture, as well as learning and memory – with locomotor activity, feeding, and social behavior being the most frequently examined parameters. Locomotor activity stands out in particular. While defecation and urination were already examined in the 1970s, wheel running, grooming, vocalization, sleep behavior, and spatial preference were studied in the home cage for the first time in the 1980s. Observations of burrowing and nesting, or behaviors (in the home cage) related to anxiety as well as human disorders like depression and schizophrenia were first reported in the 1990s. Sniffing as well as motor and sensory functions became of interest in the 2000s. Seizures were monitored in the home cage for the first time in the 2010s.

**Figure 8.**
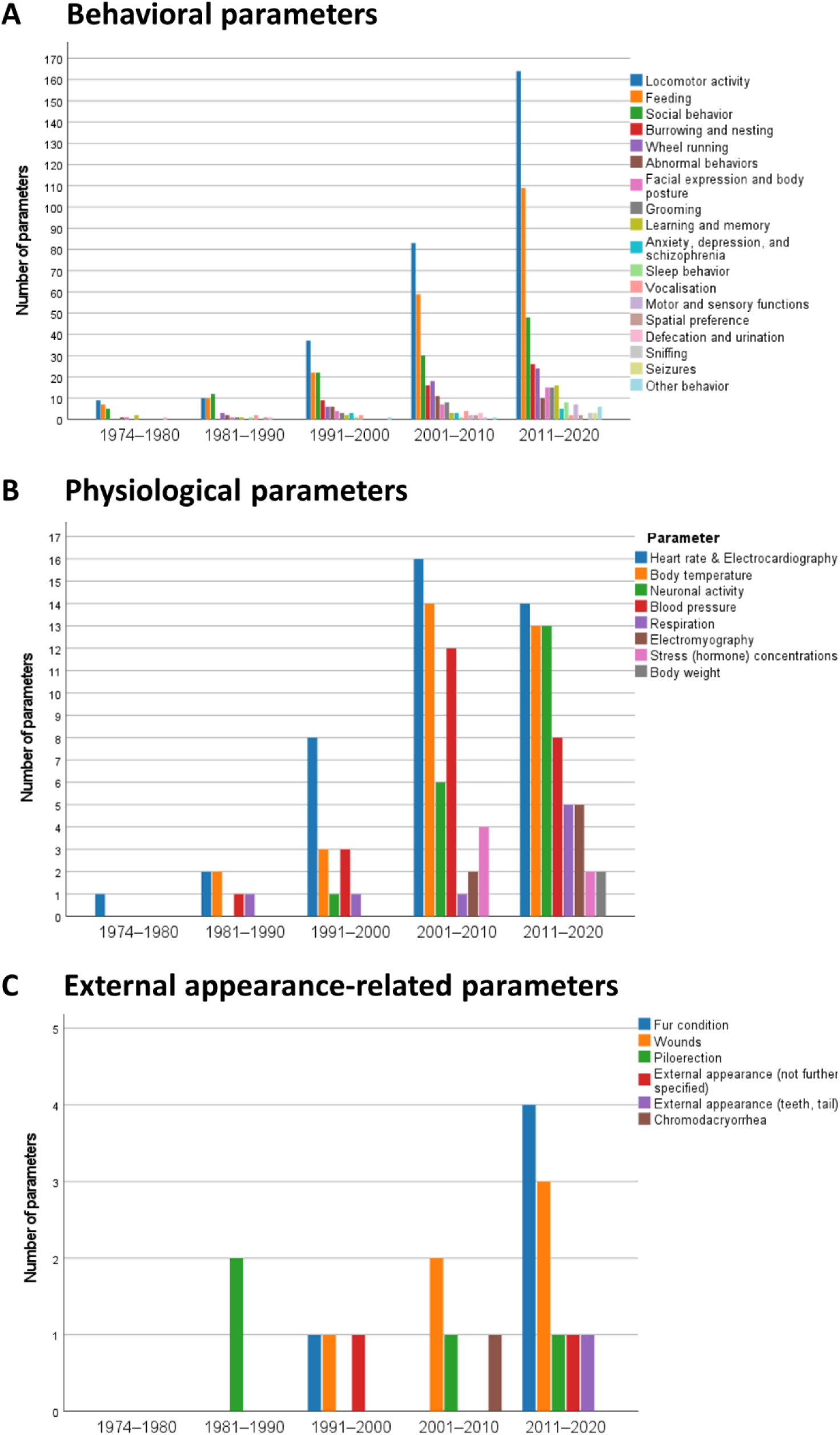
Historical change in monitoring of (A) behavioral parameters, (B) physiological parameters, and (C) external appearance-related parameters in mice and rats in their home cages. More than one application (*i.e.*, parameter) could be indicated in a publication. Other behaviors (each n = 1): colonic contractility, sneezing, sign of pain or distress (not further specified), clinical signs (not further specified), curiosity/alertness, champing behavior, twitches, behavior. Note the differences in the scaling of the y-axis.

Among the physiological parameters (Figure 8B), heart rate and/or electrocardiography were the most popular and were reported in all periods. The analysis of body temperature in the home cage became relevant in the 1980s. Its use has increased over time. It was the second most frequently examined physiological parameter in the 2000s and 2010s. Blood pressure and respiration have been measured in the home cage since 1983, neuronal activity (including electroencephalograms from implanted electrodes) since 1998, and body weight since 2017. The analysis of electromyography and hormone concentrations in the home cage were first reported between 2001 and 2010; but even later, they were investigated in a few studies only. In 2001–2010, blood pressure was the third most popular physiological parameters; in 2011–2020, neuronal activity was examined more often than blood pressure.

External appearance-related parameters have been investigated since 1988 but very few studies focus on these aspects as readouts (Figure 8C).

## Discussion

In the present systematic review, based on 521 references retrieved through PubMed and Web of Science (until Feb 2021), we studied yearly changes in HCM of laboratory mice and rats between 1974–2020. We found an increase in the use of HCM over time. The number of studies using animals of both sexes has grown, as has the study of specific disease models since the last decade(s). Over time, novel HCM techniques have been introduced and the degree of automatization has increased, allowing monitoring of more challenging animal-based parameters in the home cage. Moreover, longer housing in HCM systems with continuous monitoring, also under social housing conditions, are some of the key developments we were able to note with respect to the monitoring of mice and rats in their home cages.

### Home cage definition

The home cage definition played a central role for the present systematic review since references were only included if the following criteria were met: A home cage must allow animals to be housed permanently in their familiar social structure. We considered it important that the structure should not be changed for monitoring the parameters of interest, since this could influence the data and compromise animal welfare [16, 19]. However, animals that were kept individually throughout the process of monitoring also met our home cage definition. This definition excluded all studies in which group-housed animals were separated for the purpose of testing.

The definition of a home cage varies in the literature and may change over time with new scientific findings and ethical norms. For instance, the definition by Baran *et al.* focused on the time the animals spent in the “cages […] where animals are housed for the majority of their lifetime in the vivarium”, *i.e.*, their home cage [26]. In contrast, the definition of the present systematic review concentrated on the social structure.

To harmonize the definition of a home cage, experts in the field of HCM recently made efforts to develop a more detailed definition: www.cost-teatime.org/about/hcm-definition. This definition is structured as an olog (*i.e.*, a categorical framework for knowledge representation) [61]. It can be used for checking whether a system fulfills the criteria of HCM and it may serve as a future reporting guideline for HCM systems. However, it must be noted that the HCM definition for this systematic review was developed earlier and could not be updated after registration of the systematic review protocol.

Since the term “home cage” was part of the search string, publications that did not explicitly focus on HCM and therefore did not use this term may have not been identified by our search strategy. This includes studies where small rodents may have been monitored in their home cage.

### Strict exclusion criteria

Besides the home cage definition, the inclusion and exclusion criteria limited the number of studies that were considered. Studies involving only calorimetric measurements in the home cage were excluded since the animals usually must be transferred to a metabolic cage where they are individually housed for a short term (*i.e.*, they are separated from their social group). Due to this exclusion criteria, only those metabolic studies were included in which also other parameters (*e.g.*, locomotor activity) than metabolic outcomes were investigated.

### Publication rate

The absolute number of publications in which mice and/or rats were monitored in their home cages appeared to increase from 1974 to 2020, especially around 2005 and in the following years. Around 2005, instrumental “home cage-like” test cages were brought to the market [6, 26, 62]. A few years later, home cages with capacitance sensors, RFID and/or video tracking were developed [26]. The results of our systematic review demonstrated that custom-built systems were frequently used until the 1990s. Since the 2000s, there has been a dominance of commercial HCM systems. The high proportion of commercially available HCM systems used in recent years suggested that researchers increasingly monitored animals in their home cages since HCM systems were available on the market and more easily accessible. However, custom-built set-ups remained important tools, indicating that the HCM field is still in its early development and commercial systems currently cannot fully satisfy the demand of the user community.

In relation to the total number of PubMed-listed studies including mice or rats, the percentage of HCM publications increased over time as did the overall annual publication growth rate in Life Science, which is 5 % according to Bornemann *et al.* [63].

### Sex bias

We found a strong male-bias for both mice and rats, as reported previously for non-human mammals in other fields of biomedical research [64]. Beery and Zucker evaluated journal articles from 2009 and found a male skew in “neuroscience, physiology, pharmacology, endocrinology, and zoology” [64]. Flórez-Vargas *et al.* revealed a strong male skew in some mouse models, such as cardiovascular disease models, between 2001 and 2014 [65]. Male animals were often preferred by researchers intending to reduce potential higher data variability related the different phases of the estrous cycle in females [64]. However, the evidence for this is contentious [66–70]. A female-bias was found in reproduction and immunology [64], and infectious disease mouse models [65]. Approximately 15 % of studies published in the Journal of Physiology (London) and the Journal of Pharmacology and Experimental Therapeutics between 1909 and 2009 used non-human mammals of both sexes [64]. This percentage was a little higher (24 % over all years) in the studies included in our systematic review and, interestingly, increased since the 2010s (29 %) – probably due to a growing awareness of the consequences of a sex bias in animal-based research [71]. Finally, we would like to note the considerable advantages of testing both male and female rodents in basic science studies to better explore diseases which display variable prevalence in males and females.

### Social housing conditions

Our systematic review revealed that male mice and rats of both sexes were individually housed in the majority of studies, although individual housing can impair the emotional state of the animals. In mice, social isolation is known to increase anxiety-related behavior and depressive states [72, 73], to elevate corticosterone concentrations and reduce levels of brain-derived neurotrophic factor (BDNF) [74]. Moreover, social deprivation was shown to affect how mice react to a stressor [75]. Social deprivation is associated with welfare concerns and can result in changes in behavioral, physiological, and neurochemical parameters, metabolism, brain structures and processes [76–78]. In neurodegenerative mouse models, single housing can affect the disease progression [79]. According to our systematic review, the number of studies in which laboratory rodents were individually housed has slightly decreased in the 2000s and 2010s when compared to the previous two decades. This may also be due to the technical progress in the last decades and availability of HCM systems allowing group housing. However, because of inter-individual aggression with the onset of sexual maturity, male mice are often separated from their same-sex cage mates to avoid stress, and injuries [80–82]. It is worth noting that aggressive behavior in group housed male mice seems also to be affected by the group and cage size [83, 84]. In the studies included in our systematic review, female mice, in contrast to male mice, were more likely to be kept in groups. This may be explained by the feasibility of group housing for females. Female mice usually live in polygamous family groups in nature [85].

### Duration of housing and monitoring in the home cage

In most studies included in our review, the laboratory rodents were kept for periods of 2 days to 3 months in the HCM systems. It should be noted that data generated from animals that were only kept in the system for a few days before measurement may be biased by the novel environment and/or handling. If animals are transferred from another cage to a HCM system, it can take them a while to adapt to this novel environment. The transfer to a HCM system is comparable to a cage change. The olfactory cues as well as the familiar visual and tactile environments are removed and replaced by a clean cage, bedding, nest material, and enrichment items, which is potentially stressful for animals that strongly rely on scent for communication [86]. This can influence behavioral and physiological parameter [26, 46, 87], such as sleep [88], activity patterns [46], breathing rate [87, 89], heart rate and mean arterial blood pressure [90, 91], as well as corticosterone concentrations [92]. Since the animals are usually handled when cages are changed, the effects listed above may not only be due to a novel environment but can also be associated with the handling techniques used [18, 93].

Since habituation times are rarely reported, we refrained from extracting the time the animals were allowed to acclimatize to the respective HCM. Instead, data on the duration of measurement using the HCM techniques was collected. HCM techniques were mainly applied for short-term periods (*i.e.*, 1–28 days). The number of techniques being applied for intermediate length periods (*i.e.,* 4–12 weeks) slightly increased over time in the years between 2011 and 2020, however, 2–7 days remained the most frequent duration of measurement. Only in rare cases, HCM techniques were applied for long-term periods (*i.e.,* more than 3 months) meaning only few longitudinal studies were carried out. However, the number of studies involving measurements during the entire time in which the animals were kept in the HCM systems slightly increased over the years.

### Towards 24/7 automated home cage monitoring

HCM techniques were increasingly used for continuous data recording. Our systematic review provides evidence that the development towards 24/7 surveillance has gone hand in hand with automatization. Since the 2000s, a majority of applied HCM techniques collect data automatically, and measurements have been recorded for 24 hours a day. Long-term automated 24/7 surveillance of the animals in their home cage will allow for unbiased measurements. This is a major advance for phenotyping, monitoring of effect size of interventions, and animal welfare.

24-hour HCM takes into account the normal circadian rhythm of behavioral and physiological parameters and will reveal any phase shift relative to the light-dark-cycle (for review, see [94]). When nocturnal animals such as mice or rats are monitored for a few hours only and then often during the light phase, valuable information may be missed, as Eikelboom and Lattanzio showed for running wheel activity [15, 95].

Automatization allows for monitoring of animals without the presence of an experimenter. Prey animals such as mice and rats may show altered behavior and hide for instance pain, suffering, or distress [96]. Therefore, live cage-side health checks may not detect impaired well-being. Automated systems can thus be an important supplement to professional visual inspection of animal health and well-being. However, in our view, the cage-side checks performed by the animal care staff should never fully be replaced since there is always the risk of technical failures. It is widely discussed that an experimenter can influence animal-based parameters [97]. For example, the odor of male experimenters was shown to cause stress and stress-induced analgesia in mice and rats [98]. However, a recent multi-laboratory study demonstrated that experimenters caused less data variation than the different laboratories [99].

In contrast to manual evaluation, automatization allows for more unbiased and data driven assessment of parameters, which can save labor time once an automated method is established.

### Use of novel techniques allows for monitoring sophisticated parameters

Between 1974 and 1980, besides the manual evaluation of videos or live observations, beam based tracking [100, 101], telemetry [39], and electronic sensors and transducers [102, 103] were the first techniques applied for HCM. Interestingly, manual evaluation is still relevant in recent times and was the most frequently used technique in all time periods. The role of beam-based tracking, telemetry, as well as RFID systems became more and more important over time. Moreover, visual object tracking (video) has experienced a major boost (*i.e.*, an increase in use) since the 2010s. As data processing technology has advanced, an increased number of data parameters can be collected in the home cage.

In the 2010s, there was a shift in technology with non-invasive methods (*i.e.,* beam-based tracking and visual object tracking) overtaking invasive methods (*i.e.,* telemetry), which may indicate a shift towards improved animal welfare. The use of implanted telemetry devices compromises animal welfare. Thus, in the interest of animal welfare, non-invasive techniques should be preferred over those that are associated with a burden for the animals [104]. Although not all parameters measured by telemetry can be analyzed using beam-based tracking and visual object tracking, activity can be monitored using non-invasive methods. Moreover, non-invasive alternatives such as jacketed telemetry could be applied for assessing respiration, collecting an electrocardiogram, or measuring activity [105].

Most studies conducted so far with HCM (69 %) used only one technique. However, studies collecting a larger set of parameters or extracted data from individuals among group-housed animals usually deployed multiple techniques to increase the versatility of the HCM system. The present systematic review revealed an increase in the number of HCM studies involving disease models, which may be attributed to the technical progress to examine more and more parameters in the home cage and the demand to characterize a rapidly growing number of genetic mutants since the 2000s.

External appearance-related parameters (*e.g.*, body condition score, wounds, fur condition, piloerection, chromodacryorrhea) have only been rarely observed in the home cage yet although they are key for tracking animals’ welfare status. We envisage that machine learning approaches will play a central role in HCM data processing and in the future may also enable automatic monitoring of external appearance-related parameters. Unsupervised and supervised machine learning can be used to automatically analyze video frames recorded in the home cage [106–108].

All in all, our systematic review revealed that HCM in mice and rats has gone through a considerable development and as these instruments have become more comprehensive, easy to apply, and scalable, the use of HCM has increased. There is a slight trend towards recordings covering 24 hours over intermediate length periods as the HCM techniques are refined and become more and more automated and application of HCM is spreading to a wider range of study types. A considerable fraction of the HCM systems is still custom-built, but commercial “key ready” solutions are increasingly available and since the 2000s more than 50 % of the HCM studies used commercial products.

Although, manual observations made live or from videos remain key technical solutions for HCM, this review indicates that a number of alternative, high-throughput, and man-power saving techniques have been introduced.

Future inventions will pave the way for continuous, non-invasive rodent monitoring in longitudinal studies. Storage and analysis of the large amounts of data generated by automated HCM are a bottleneck that needs to be addressed [26]. Moreover, the financial aspects may be a hurdle in implementing commercial systems.

## Acknowledgement

This article is based upon work from COST Action “Improving biomedical research by automated behaviour monitoring in the animal home-cage” (TEATIME; CA20135; cost-teatime.org) supported by COST (European Cooperation in Science and Technology).

## Supporting information

**S1 Supporting Information. Data extraction template.**

**S2 Supporting Information. Objective, Materials & Methods. S3 Supporting Information. Mouse and rat strains.**

**S4 Supporting Information. Home cage monitoring system.**

**S5 Supporting Information. Behavioral, physiological, and external appearance-related parameters.**

**S6 Data. Objective, Materials & Methods.**

