## Supplementary material for "Development and Application of Home Cage Monitoring in Laboratory Mice and Rats: a Systematic Review": S1 Supporting information

**S1 Supporting Information – Data extraction template**

### Covidence – Data extraction template

*Note: Please make sure not to use commas in the free text entries.*

#### 1. General information

##### 1.1 Journal

*Copy the name of the journal from the PDF (full name, no abbreviation).*

#### 2. Animal model

##### 2.1 Species

- ○ Mouse
- ○ Rat
- ○ Both mouse and rat
- ○ N/A

##### 2.2 Strain

*For non-genetically modified animals, enter the strain; for genetically modified animals, indicate the background strain. If "strain" is not applicable, please also select "other" and specify.*

- ○ C57BL/6
- ○ BALB/c
- ○ DBA/2
- ○ Swiss Webster
- ○ NMRI
- ○ CD-1
- ○ Long Evans
- ○ Wistar
- ○ N/A
- ○ More than one strain
- ○ Other: ________________

##### 2.3 Please specify "more than one strain".

*If "more than one strain" was used, copy the species from the list above and paste them in the field "other". Use "AND" between several entries, e.g. C57BL/6 AND CD-1. If "more than one strain" is not applicable, select "N/A".*

- ○ N/A
- ○ Other: ________________

##### 2.4 Sex

- ○ Male
- ○ Female
- ○ Both
- ○ N/A

**2.5 Disease model**

*A disease model has to 1) be a disease according to ICD-11 (icd.who.int/browse11/l-m/en) and 2) the authors themselves have to define it as a disease model (unless it is very obviously a disease model).*

- No disease model
- Infectious or parasitic diseases
- Neoplasms
- Diseases of the blood or blood-forming organs
- Diseases of the immune system
- Endocrine, nutritional or metabolic diseases
- Mental, behavioral or neurodevelopmental disorders
- Sleep-wake disorders
- Diseases of the nervous system
- Diseases of the visual system
- Diseases of the ear or mastoid process
- Diseases of the circulatory system
- Diseases of the respiratory system
- Diseases of the digestive system
- Diseases of the skin
- Diseases of the musculoskeletal system or connective tissue
- Diseases of the genitourinary system
- More than one disease model
- Other: ________________

**2.6 Please specify "more than one disease model".**

*If "more than one disease model" was used, copy the disease models from the list above and paste them in the field "Other". Use "AND" between several entries. If "more than one disease model" is not applicable, select "N/A".*

- N/A
- Other: ________________

#### 3. Home cage system

**3.1 Which home cage system was used?**

*If a commercially available system was used, indicate its name. "Other" can be other commercially available systems. A "custom-built" system is defined as 1) not commercially available for which the authors may provide construction plans and/or software and 2) to which the authors have given a particular name, e.g. Live Mouse Tracker. Choose "N/A" if neither a commercially available nor a custom-built system was used, e.g. the sole use of cameras.*

- Activmetre (Bioseb)
- AfaSci
- Any-maze Cage (Stoelting)
- DVC® Tecniplast
- DSI’s Telemetry Devices
- HCA (Actual analytics)
- HomeCageScan (Cleversys)
- Infrared Motion Detector (Starr Life Tecnologies)

- Intellicage (TSE)
- Kinder Scientific
- Laboras (Metris)
- Phenotyper Noldus
- Photobeam Activity System (San Diego Instruments)
- Smart Cage (Omnitech Electronics)
- PhenoMaster (TSE)
- Uga Basile
- Videotrack (Viewpoint)
- Custom-built
- N/A
- More than one system
- Other: ________________

**3.2 Please specify "more than one system".**
*If "more than one system" was used, copy the systems from the list above and paste them in the field "Other". Use "AND" between several entries. If "more than one system" is not applicable, select "N/A".*
- N/A
- Other: ________________

**3.3 Number of animals housed in the home cage system**
*Please fill in the number of animals that are kept in a home cage system (this does not have to be the total number of animals used in the study but only the number of animals living together in a system during the monitoring period). If the study consists of several substudies with different numbers of animals housed in the home cage system, fill in the maximum number.*
- 1
- 2
- 3
- 4
- 5
- 6
- 7
- 8
- 9
- 10
- More than 10
- N/A

**3.4 Duration spent in the home cage system**
*Please fill in the total number of days the animals were housed in the home cage system (including periods with and without monitoring). If the study consists of several substudies with a variation in this parameter, fill in the maximum duration. If indicated as weeks or months, calculate as follows:*
- 1 week = 7 days; 1 month = 30 days
- 1 day
- 2-7 days
- 8-14 days (1-2 weeks)

- 15-28 days (2-4 weeks)
- 29-84 days (1-3 months)
- 85-168 days (3-6 months)
- 169-336 days (6-12 months)
- 337 and more days (> 1 year)
- N/A

#### 4. Home cage monitoring techniques

**4.1 How many home cage monitoring techniques were used in the study?**

- 1
- 2
- 3
- more than 3
- animals were not monitored in the home cage but in the testing apparatus connected to the home
- cage
- N/A

***The following questions were repeated three times, i.e., for three techniques – 4.2, 4.3, and 4.4.***

**4.2 Home cage monitoring technique #1 (Fill in the techniques in alphabetical order, e.g. #1 RFID system, #2 thermal imaging).**

**4.2.1 Which is home cage monitoring technique #1 (1st in alphabetical order)?**

- Body weight sensors
- Electromagnetic detection
- Infrared based system
- Manual (live) monitoring
- Microwave based system
- RFID system
- Short-wavelength infrared region (SWIR) imaging
- Telemetry
- Thermal imaging
- Vibration and force senstive plates
- Vibration sensitive plate
- Video (RGB) – manual evaluation
- Video (RGB) tracking system
- Video (RGB-D) tracking system
- N/A
- Other: ________________

**4.2.2 Which parameters were investigated using technique #1?**

- Abnormal behaviors
- Anxiety, depression, and schizophrenia
- Body Condition Score
- Body temperature
- Body weight

- Burrowing and nesting
- Electroencephalogram
- Facial expression and body posture
- Food and water intake
- Fur condition
- Heart rate
- Learning and memory
- Locomotor activity
- Motor and sensory functions
- Pilorection
- Respiration
- Social behavior
- Stress hormones
- Vocalisation
- Wheel running
- Wounds
- N/A
- More than one parameter
- Other: ________________

**4.2.3 Please specify "more than one parameter" that were investigated using technique #1.**
*If "more than one parameter" was used, copy the parameter from the list above and paste them in the field "Other". Use "AND" between several entries. If "more than one parameter" is not applicable, select "N/A".*

- N/A
- Other: ________________

**4.2.4 Degree of automatization of technique #1**
*1) Manual: without technological aids (technological aids do not include cameras used for manual monitoring), e. g. manual (live) monitoring, video (RGB) – manual evaluation. 2) Automatic: with technological aids, e. g. video (RGB) tracking system.*

- Manual
- Automatic
- N/A

**4.2.5 Monitoring of individuals or groups using technique #1**
*Please indicate whether values of individuals (i.e. each animal of a group) or values of groups (i.e. the animal group as a whole) were determined.*

- Individuals
- Groups
- N/A

**4.2.6 Duration of measurement per day using technique #1**
*Please also include the hours that were excluded from the analysis (if reported).*

- 24 hours
- More than 12 and less than 24 hours
- Less than 12 hours

- N/A

**4.2.7 Overall duration of measurement using technique #1**

*Please fill in the number of days the animals were monitored in the home cage system. This is NOT the sum of all measurement hours but the number of days on which measurements were performed. If the study consists of several substudies with a variation in this parameter, fill in the maximum duration. If indicated as weeks or months, calculate as follows: 1 week = 7 days; 1 month = 30 days*

- 1 day
- 2-7 days
- 8-14 days (1 to 2 weeks)
- 15-28 days (2-4 weeks)
- 29-84 days (1-3 months)
- 85-168 days (3-6 months)
- 169-336 days (6-12 months)
- 337 and more days (> 1 year)
- N/A
