## Supplementary material for "Development and Application of Home Cage Monitoring in Laboratory Mice and Rats: a Systematic Review": S2 Supporting information

### S2 Supporting Information – Objective, Materials & Methods

#### Objective

**Methods/concept of interest:** We investigated home cage monitoring techniques that were used to analyze behavioral, physiological, and/or external appearance-related parameters in mice and rats. A home cage was defined as any cage in which the animals could potentially be housed permanently in their familiar social structure (*i.e.,* group or single housing). A testing apparatus could be connected to the home cage allowing the animals to voluntarily enter it. In this scenario, the animals may also separate themselves from the group when entering the testing apparatus. The home cage could be permanently or temporarily divided so that the animals are separated from each other by a cage divider (*e.g.,* a grid). Our home cage definition excluded any cage system that required the separation of group-housed mice or rats for the duration of monitoring. In contrast, if an animal was permanently kept socially isolated in a cage system, the home cage definition was met.

**Species:** All primary studies on mice (Genus: *Mus*) or rats (Genus: *Rattus*) of any age were included.

**Outcome measures:** The outcome measures were techniques and applications for home cage monitoring and the degree of automatization.

**Research question:** How have techniques and applications for home cage monitoring of laboratory mice and rats developed over time? Has the degree of automatization for monitoring behavioral, physiological, and external appearance-related parameters changed?

### S2 Supporting Information – Objective, Materials & Methods

#### Materials & methods

##### Study selection

Screening phases: Search results were imported into EndNote (Version X9.3.3). Duplicates were removed and the full text of articles, if available, were retrieved using the built-in features. The remaining entries were transferred from EndNote to Covidence (1) where remaining duplicates were removed and all screening phases were carried out. Papers were processed in three consecutive phases: In phase 1, titles and abstracts were screened. Thereafter, full texts were screened (phase 2) and data were extracted (phase 3) simultaneously.

Number of observers per screening phase and discrepancies: In all phases, all papers were screened by two independent reviewers and discrepancies were resolved by a third reviewer. For this, the reviewer settling discrepancies had access to the notes of the other two screeners.

Inclusion and exclusion criteria:

- Type of study
    - Inclusion criteria: primary study
    - Exclusion criteria: survey, interview, non-primary research/review, conference abstract only, not a full peer-reviewed article
- Type of animals
    - Inclusion criteria: mice (Mus), rats (Rattus)
    - Exclusion criteria: not mice (Mus) or rats (Rattus)
- Type of intervention
    - Inclusion criteria: monitoring in home cage or testing apparatuses connected to home cage
    - Exclusion criteria: monitoring not in home cage or testing apparatuses not connected to home cage
- Outcome measures
    - Inclusion criteria: Parameters investigated using a home cage monitoring technique (behavioral parameters, physiological parameters, external appearance-related parameters)
    - Exclusion criteria: calorimetry only (studies in which no other parameters than calorimetric measurements were examined in a home cage)
- Language restrictions
    - Inclusion criteria: English
    - Exclusion criteria: Not English language

Important to note is that the resident intruder test was not an exclusion criterion although the familiar social structure of the mice in the home cage is changed for a short period of time during this test.

#### S2 Supporting Information – Objective, Materials & Methods

##### Study characteristics to be extracted

The data extraction template used in Covidence can be found in Supplemental Material 1. In brief, the following data were of interest:

- Study meta-data: First author, year, title, and journal were extracted.
- Study design characteristics: If a particular home cage system was used, its name was noted. The number of animals housed in the home cage system and the duration spent in the home cage system were extracted.
- Animal model characteristics: Species, strain/stock, sex, and disease model were extracted. For genetically modified animals, the background strain was extracted. C57BL/6J and C57BL/6N were subsumed under C57BL/6.
- Types of methods: The monitoring techniques as well as the degree of automatization were extracted. The latter referred to the way parameters were recorded using a particular technique: manually (*i.e.*, without technological aids, with the exception for the use of cameras where data was recorded manually from the recordings) or automatically (*i.e.*, with technological aids like a video tracking system). Moreover, we extracted whether individuals or groups of animals were studied.
- Outcomes measures: Parameters that were recorded using a home cage monitoring technique were extracted. In addition, the duration of measurement per day and the overall duration of measurement were extracted.

The following home cage monitoring techniques were defined. If a technique was not listed, it was indicated as free-text response.

- Body weight sensors
- Infrared based system (i.e., one or even more rows of infrared beams surrounding the cage)
- Manual (live) monitoring (i.e., the experimenter monitors the animal(s) in front of the cage, through a window or using a camera and assesses their behavior (live))
- Microwave based system (i.e., radar can be used to detect motions)
- RFID system
- Short-wavelength infrared region (SWIR) imaging
- Telemetry
- Thermal imaging
- Vibration and force sensitive plates
- Vibration sensitive plate
- Video (RGB) – manual evaluation (i.e., videos from RGB (red-green-blue) cameras are evaluated by hand, the experimenters watch the videos and assess the behavior of the animals)
- Video (RGB) tracking system (i.e., videos from RGB cameras are evaluated using a software)
- Video (RGB-D) tracking system (i.e., videos from RGB-D (red-green-blue with depth) cameras are used for behavioral assessment)

#### S2 Supporting Information – Objective, Materials & Methods

For data analysis, these techniques were grouped as follows:

- Audio recording
- Beam based tracking: infrared based system, laser beam based system, photointerrupter
- Drinkometer/Lickometer: drinkometer, lickometer, volumetric drinking tubes
- Electronic sensors and transducers: microwave based system, electromagnetic detection, electrical conductance, piezoelectric sensor, capacitance sensor, electronic strain gauge-based load cell
- Manual evaluation: manual (live) monitoring, video (RGB) – manual evaluation, infrared camera (manual)
- RFID
- Running wheel (counter)
- Telemetry
- Tissue sampling: microdialysis, automated blood sampling, jugular vein catheter
- Vibration, force and weight sensitive tracking: vibration and force sensitive plates, vibration sensitive plate, body weight sensors, scale, weight sensors, gravimetric analysis
- Visual object tracking (video): video (RGB) tracking system, video (RGB-D) tracking system, motion-activated cameras, infrared camera (automatic)
- Other: fiber photometry system, brain imaging cameras, automatic food dispenser, lever, cardiotachometer, flowmeter circuit, infrared thermometer, thermal imaging, impedance pneumography

Only parameters that were monitored in the home cage of the animals were extracted from the studies. The behavioral, physiological, and external appearance related parameters were defined, as follows below. If a parameter was not listed, it was indicated as free-text response.

- Behavioral parameters
    - Abnormal behaviors: e.g., infanticide, barbering, stereotypy
    - Anxiety, depression, and schizophrenia: e.g., defensive burying, Vogel conflict test, Geller-Seifter conflict test, sucrose preference test, prepulse inhibition, sensitization to psychostimulants, latent inhibition
    - Burrowing and nesting
    - Facial expression and body posture: pain faces such as Mouse and Rat Grimace Scale, facial expressions displayed in other states than pain, body posture associated with pain or any emotional state of an individual (excl. social interaction)
    - Feeding (drinking, food): food intake, water intake, intake of other liquids, two bottle choice test
    - Learning and memory: e.g., contextual and cued fear conditioning, social recognition, object recognition, other tests associated with exploration, social transmission of food preference, passive avoidance, active avoidance, operant nose-poke reinforcement, eyeblink conditioning, conditioned taste preference, conditioned taste aversion, conditioned place preference
    - Locomotor activity
    - Motor and sensory functions: e.g., grip gauge, footprint analysis, balance beam, gait, ataxia, hot plate (nociception)

    - Social behavior: e.g., social interaction/preference, social grooming, social dominance, aggression, sexual behavior, maternal behavior, juvenile play
    - Vocalization
    - Wheel running

- Physiological parameters
    - Body temperature
    - Body weight
    - Electroencephalography
    - Heart rate
    - Respiration
    - Stress hormones

- External appearance
    - Body Condition Score
    - Fur condition
    - Pilorection
    - Wounds
