## Supplementary material for "Development and Application of Home Cage Monitoring in Laboratory Mice and Rats: a Systematic Review": S3 Supporting information

### S3 Supporting Information – Mouse and rat strains

After all data were extracted from the included references in phase 3 and a consensus reviewer clarified any conflicts, all entries for “strain” were double-checked by an additional reviewer (OK or EG) and corrected if necessary.

Following corrections were made:

- C57BL/6J and C57BL/6N were subsumed under C57BL/6, as requested in the instructions of the systematic review.
- 129S, 129S6, and SvEvTac were subsumed under 129.
- ICR and CD-1 were subsumed under CD-1
- Holtzman and Sprague Dawley were subsumed under Sprague Dawley.
- We differentiated between Wistar and Wistar Kyoto.
- For the genetically modified strains, the background strain was indicated. If they were on a mixed background, this was listed as "mixed background."
- If the strain was not further specified in the reference and only “hooded” or “albino” was indicated, the answer for strains was changed to “N/A” (not applicable).

### S3 Supporting Information – Mouse and rat strains

| Mouse strain | Number of publications |
|---|---|
| C57BL/6 | 192 |
| BALB/c | 31 |
| CD-1 | 26 |
| Mixed background | 24 |
| DBA/2 | 21 |
| 129 | 16 |
| FVB | 9 |
| Swiss Webster | 8 |
| CBA | 6 |
| C3H | 6 |
| NOD | 5 |
| A/J | 4 |
| BTBR | 4 |
| AKR | 2 |
| Binghamton heterogeneous | 2 |
| CAST | 2 |
| NMRI | 2 |
| NZB | 2 |
| TO | 2 |
| N/A | 2 |
| 129S | 1 |
| BKW | 1 |
| C57BL/10 | 1 |
| CB17 | 1 |
| CF1 | 1 |
| Collaborative Cross | 1 |
| db/db | 1 |
| DD | 1 |
| LACA | 1 |
| MF1 | 1 |
| NIH | 1 |
| NIH Swiss | 1 |
| NSY | 1 |
| NZO | 1 |
| OF-1 | 1 |
| PWK | 1 |
| SCID | 1 |
| SKH | 1 |
| Swiss SE | 1 |
| SWR | 1 |
| Wild mouse *Mus musculus domesticus* | 1 |
| WSB | 1 |
| WSC | 1 |
| WSP | 1 |

More than one strain could be used per study.

### S3 Supporting Information – Mouse and rat strains

| Rat strain | Number of publications |
|---:|---:|
| Sprague Dawley | 115 |
| Wistar | 74 |
| Long Evans | 34 |
| Lister hooded | 9 |
| Fischer 344 | 7 |
| SHR | 5 |
| Lewis | 3 |
| Brown Norway | 3 |
| NIH | 3 |
| Wistar Kyoto | 3 |
| Black hooded | 2 |
| Hooded | 2 |
| Tryon maze dull | 2 |
| Alcohol-preferring P rat | 2 |
| N/A | 3 |
| BDIX | 1 |
| Berkeley S1 | 1 |
| Chester Beatty | 1 |
| Fawn Hooded | 1 |
| Flinders sensitive line | 1 |
| MNRA | 1 |
| MR | 1 |
| RHA-I | 1 |
| RLA-I | 1 |
| Sabra | 1 |
| Sherman | 1 |

More than one strain could be used per study.
