## Supplementary material for "Development and Application of Home Cage Monitoring in Laboratory Mice and Rats: a Systematic Review": S4 Supporting information

#### S4 Supporting Information – Home cage monitoring system

The most frequently used commercially available HCM systems were the Data Sciences International telemetry devices (DSI, USA; n = 48). Other telemetry devices were produced by Starr Life Sciences (USA, E-Mitter Telemetry Implants, n = 7), Mini-Mitter (USA, n = 2), and Transoma Medical (USA, telemetry probe, n = 1).

A range of running/activity wheels from different companies have been used: Lafayette Instruments (USA, n = 4), Mini-Mitter (USA, n = 2), MED Associates (USA, n = 2), ClockLab, Actimetrics (USA, n = 2), TSE (USA/Germany; n = 2), Wahmann Mfg. Co., Model LC-34, n = 1), Tecniplast (Italy, n = 1), Shinano Ltd. (Japan, n = 1), Respironics (USA, n = 1). In one study, the manufacturer was not indicated.

Other HCM systems found in publications were the Phenotyper (Noldus, the Netherlands; n = 26), HomeCageScan (Cleversys, USA; n = 10), IntelliCage (TSE, USA/Germany; n = 9), PhenoMaster (TSE, USA/Germany; n = 7), InfraMot (TSE, USA/Germany; n = 6), DVC (Tecniplast, Italy; n = 5), HCA (Actual analytics, United Kingdom; n = 5), Laboras (Metris, the Netherlands; n = 4), EthoVision (Noldus, the Netherlands; n = 4), Activity monitor (AccuScan Instruments, USA; n = 4), Opto-M3 Dual Axis System (Columbus Instruments, USA; n = 4), Operant chambers (MED Associates, USA; n = 3), ActiviScope (NewBehavior, Switzerland; n = 3), 24 channel activity monitoring system (O'Hara & Co., Tokyo, Japan; n = 3), Home Cage Activity System (Coulbourn Instruments, Allentown, PA, USA; n = 3), operant chambers (Coulbourn Instruments, USA; n = 2), MicroMax Activity monitor (AccuScan Instruments, USA; n = 2), Animex activity meter (LKB Instruments, Sweden; n = 2), and Infrared Motion Detector (Starr Life Sciences, USA; n = 2).

#### S4 Supporting Information – Home cage monitoring system

| Home cage monitoring systems | Number of publications | Species (multiple answers possible) | Strain (multiple answers possible) | Sex | Number of animals housed in the home cage system | Duration spent in the home cage system |
|---|---|---|---|---|---|---|
| N/A | 209 | | | | | |
| More than one system | 28 | | | | | |
| Custom-built | 126 | | | | | |
| **Telemetry** | | | | | | |
| Telemetry Devices (DSI, USA) | 48 (40) | Mouse (n=28)<br>Rat (n=20) | C57BL/6 (n=17)<br>CD-1 (n=4)<br>BALB/c (n=3)<br>FVB (n=2)<br>DBA/2 (n=1)<br>Swiss SE (n=1)<br>Mixed background (n=2)<br>Sprague Dawley (n=10)<br>Wistar (n=9)<br>SHR (n=3)<br>BDIX (n=1)<br>Brown Norway (n=1)<br>Fischer 344 (n=1)<br>Wistar Kyoto (n=1) | Male (n=36)<br>Female (n=7)<br>Both (n=4)<br>N/A (n=1) | 1 (n=34)<br>2 (n=5)<br>3 (n=2)<br>4 (n=1)<br>N/A (n=6) | 1 day (n=1)<br>2–7 days (n=5)<br>8-14 days (1–2 weeks) (n=8)<br>15-28 days (2–4 weeks) (n=12)<br>29-84 days (1–3 months) (n=7)<br>85-168 days (3–6 months) (n=4)<br>N/A (n=11) |
| E-Mitter Telemetry Implants (Starr Life Sciences, USA) | 7 | Mouse (n=5)<br>Rat (n=2) | C57BL/6 (n=3)<br>DBA/2 (n=1)<br>NIH Swiss (n=1)<br>SHR (n=1)<br>Sprague Dawley (n=1)<br>Wistar Kyoto (n=1) | Male (n=5)<br>Female (n=2) | 1 (n=4)<br>2 (n=1)<br>N/A (n =2) | 2–7 days (n=1)<br>8-14 days (1–2 weeks) (n=3)<br>15-28 days (2–4 weeks) (n=2)<br>29-84 days (1–3 months) (n=1) |
| Telemetry probe (Transoma Medical, USA) | 1 | Rat (n=1) | Sprague Dawley (n=1) | Male (n=1) | 1 (n=1) | 2–7 days (n=1) |
| Telemetry (Mini-Mitter, USA) | 2 | Mouse (n=2) | C57BL/6 (n=1)<br>AKR (n=1)<br>SWR (n=1) | Male (n=2) | 1 (n=2) | N/A (n=2) |

### S4 Supporting Information – Home cage monitoring system

| | | | | | | |
|---|---|---|---|---|---|---|
| | | | db/db (n=1) | | | |
| **Wheel running** | | | | | | |
| Running wheels (Mini-Mitter, USA) | 2 | Mouse (n=2) | C57BL/6 (n=1)<br>Mixed background (n=1) | Male (n=1)<br>Both (n=1) | 1 (n=1)<br>4 (n=1) | 8–14 days (1–2 weeks) (n=1)<br>N/A (n=1) |
| Running wheels (MED Associates, USA) | 2 | Mouse (n=1)<br>Rat (n=1) | C57BL/6 (n=1)<br>Sprague Dawley (n=1) | Male (n=2) | 1 (n=2) | 8–14 days (1–2 weeks) (n=1)<br>29–84 days (1–3 months) (n=1) |
| Activity wheels (Wahmann Mfg. Co., Model LC-34) | 1 | Rat (n=1) | Wistar (n=1) | Male (n=1) | 1 (n=1) | 15–28 days (2–4 weeks) (n=1) |
| Activity wheels (Lafayette Instruments, USA) | 4 | Mouse (n=3)<br>Rat (n=1) | C57BL/6 (n=3)<br>Long Evans (n=1) | Male (n=2)<br>Both (n=2) | 1 (n=2)<br>N/A (n=2) | 15–28 days (2–4 weeks) (n=1)<br>29–84 days (1–3 months) (n=1)<br>85–168 days (3–6 months) (n=1)<br>169–336 days (6–12 months) (n=1) |
| Running wheels (manufacturer not indicated) | 1 | Rat (n=1) | Sprague Dawley (n=1) | Female (n=1) | 1 (n=1) | 85–168 days (3–6 months) (n=1) |
| Running wheels (ClockLab, Actimetrics, USA) | 2 | Mouse (n=1)<br>Rat (n=1) | Mixed background (n=1)<br>Wistar (n=1) | Male (n=1)<br>Both (n=1) | 1 (n=2) | 2–7 days (n=1)<br>29–84 days (1–3 months) (n=1) |
| Running wheels (Tecniplast, Italy) | 1 | Rat (n=1) | Wistar (n=1)<br>Sprague Dawley (n=1) | Male (n=1) | 2 (n=1) | 29–84 days (1–3 months) (n=1) |
| Running wheels (TSE, USA/Germany) | 2 | Mouse (n=2) | C57BL/6 (n=2)<br>NZB (n=1)<br>NZO (n=1) | Male (n=1)<br>N/A (n=1) | 1 (n=2) | 2–7 days (n=1)<br>N/A (n=1) |
| Running wheels (Shinano Ltd, Japan) | 1 | Rat (n=1) | Wistar (n=1) | Male (n=1) | 1 (n=1) | 8–14 days (1–2 weeks) (n=1) |
| Running Wheels (Respironics, USA) | 1 | Mouse (n=1) | CD-1 (n=1)<br>Collaborative Cross (n=1) | Both (n=1) | 1 (n=1) | 2–7 days (n =1) |

### S4 Supporting Information – Home cage monitoring system

**The following systems were listed in alphabetical order.**

| | | | | | | |
|---|---|---|---|---|---|---|
| 24 channel activity monitoring system (O'Hara & Co., Tokyo, Japan) | 3 | Mouse (n=3) | C57BL/6 (n=3) | Male (n=3) | 1 (n=2)<br>4 (n=1) | 2–7 days (n=1)<br>8–14 days (1–2 weeks) (n=1)<br>29–84 days (1–3 months) (n=1) |
| ACTIMO System (Shintechno, Japan) | 1 | Rat (n=1) | Wistar (n=1) | Male (n=1) | 1 (n=1) | 8–14 days (1–2 weeks) (n=1) |
| ActiviScope (NewBehavior, Switzerland) | 3 | Mouse (n=2)<br>Rat (n=1) | C57BL/6 (n=1)<br>Mixed background (n=1)<br>Wistar (n=1) | Male (n=2)<br>N/A (n=1) | 1 (n=3) | 15–28 days (2–4 weeks) (n=1)<br>29–84 days (1–3 months) (n=1)<br>N/A (n=1) |
| Activity cages, UMOTWin (Ellegaard systems, Denmark) | 1 | Mouse (n=1) | Mixed background (n=1) | Both (n=1) | 1 (n=1) | 2–7 days (n=1) |
| Activity monitor (AccuScan Instruments, USA) | 4 | Mouse (n=4) | C57BL/6 (n=3)<br>129 (n=1)<br>WSC (n=1)<br>WSP (n=1) | Male (n=3)<br>Female (n=1) | 1 (n=4) | 8–14 days (1–2 weeks) (n=2)<br>15–28 days (2–4 weeks) (n=1)<br>29–84 days (1–3 months) (n=1) |
| Activity sensor (Neuroscience Inc., Japan) | 1 | Mouse (n=1) | C57BL/6 (n=1)<br>BALB/c (n=1) | Male (n=1) | N/A (n=1) | 29–84 days (1–3 months) (n=1) |
| Activity sensor and food intake monitor (O'Hara & Co., Tokyo, Japan) | 1 | Rat (n=1) | Wistar (n=1) | Male (n=1) | 1 (n=1) | 29–84 days (1–3 months) (n=1) |
| Animex activity meter (LKB Instruments, Sweden) | 2 | Mouse (n=1)<br>Rat (n=1) | C57BL/6 (n=1)<br>Sprague Dawley (n=1) | Male (n=2) | 2 (n=1)<br>10 (n=1) | 15–28 days (2–4 weeks) (n=1)<br>29–84 days (1–3 months) (n=1) |
| Any-maze Cage (Stoelting, Ireland) | 1 | Mouse (n=1) | C57BL/6 (n=1) | Male (n=1) | 1 (n=1) | 8–14 days (1–2 weeks) (n=1) |
| Automated drinking & feeding monitor system (TSE, USA/Germany) | 1 | Mouse (n=1) | C57BL/6 (n=1)<br>NZB (n=1)<br>NZO (n=1) | Male (n=1) | n (n=1) | N/A (n=1) |
| BASi (West Lafayette, USA) | 1 | Rat (n=1) | Wistar (n=1) | N/A (n=1) | N/A (n=1) | 8–14 days (1–2 weeks) (n=1) |
| BioDAQ food intake monitoring system (Research Diets, USA) | 1 | Rat (n=1) | Wistar (n=1) | Female (n=1) | 1 (n=1) | N/A (n=1) |

### S4 Supporting Information – Home cage monitoring system

| | | | | | | |
|---|---|---|---|---|---|---|
| CI Multi-Device Interface Multi Device Interface MDI Software (Columbus Instruments, USA) | 1 | Mouse (n=1) | C57BL/6 (n=1) | Male (n=1) | 1 (n=1) | 15–28 days (2–4 weeks) (n=1) |
| Comprehensive Laboratory Animal Monitoring System (CLAMS, Columbus Instruments, USA | 1 | Mouse (n=1) | C57BL/6 (n=1) | Both (n=1) | 1 (n=1) | 169–336 days (6–12 months) (n=1) |
| Digiscan (AccuScan Instruments, USA) | 1 | Mouse (n=1) | BALB/c (n=1)<br>C57BL/6 (n=1)<br>DBA/2 (n=1) | Male (n=1) | 1 (n=1) | 29–84 days (1–3 months) (n=1) |
| Digital scale (EAGDCE-L, Sartorius AG, Germany) | 1 | Rat (n=1) | Sprague Dawley (n=1) | Male (n=1) | 2 (n=1) | 15–28 days (2–4 weeks) (n=1) |
| Doppler radar (Model BBL; McEwan Technologies, USA); infrared non-contact thermal imager (Flir Systems, ThermoVision A320, USA); Model 12 polygraph (Grass-Telefactor, USA) | 1 | Rat (n=1) | Wistar (n=1) | Male (n=1) | 1 (n=1) | N/A (n=1) |
| Drinkometer system, MOUSE-E-MOTION (INFRA-E-MOTION GmbH, Germany) | 1 | Mouse (n=1) | C57BL/6 (n=1) | Male (n=1) | 1 (n=1) | 29–84 days (1–3 months) (n=1) |
| DVC (Tecniplast, Italy) | 5 | Mouse (n=5) | C57BL/6 (n=4)<br>BALB/c (n=1) | Male (n=1)<br>Both (n=3)<br>N/A (n=1) | 1 (n=1)<br>2 (n=1)<br>3 (n=1)<br>4 (n=1)<br>5 (n=1) | 2–7 days (n=1)<br>29–84 days (1–3 months) (n=1)<br>85–168 days (3–6 months) (n=2)<br>337 and more days (> 1 year) (n=1) |
| EthoVision (Noldus, the Netherlands) | 4 | Mouse (n=4) | C57BL/6 (n=3)<br>Mixed background (n=1) | Male (n=2)<br>Both (n=1) | 1 (n=3)<br>N/A (n=1) | 8–14 days (1–2 weeks) (n=1)<br>N/A (n=3) |

### S4 Supporting Information – Home cage monitoring system

| | | | | | | |
|---|---|---|---|---|---|---|
| | | | | N/A (n=1) | | |
| Feeding chambers (Med Associates, USA) | 1 | Rat (n=1) | Sprague Dawley (n=1)<br>Wistar (n=1) | Male (n=1) | 2 (n=1) | 29–84 days (1–3 months) (n=1) |
| HCA (Actual analytics, United Kingdom) | 5 | Mouse (n=2)<br>Rat (n=3) | C57BL/6 (n=2)<br>C3H (n=1)<br>Sprague Dawley (n=1)<br>Wistar (n=2) | Male (n=1)<br>Both (n=3)<br>N/A (n=1) | 1 (n=1)<br>2 (n=1)<br>3 (n=1)<br>4 (n=1)<br>5 (n=1) | 2–7 days (n=1)<br>29–84 days (1–3 months) (n=1)<br>85–168 days (3–6 months) (n=2)<br>337 and more days (> 1 year) (n=1) |
| HM-2, MBRose, Denmark | 1 | Mouse (n=1) | C57BL/6 (n=1) | Male (n =1) | 4 (n=1) | 29–84 days (1–3 months) (n=1) |
| Home Cage Activity System (Coulbourn Instruments, Allentown, PA, USA) | 3 | Mouse (n=2)<br>Rat (n=1) | C57BL/6 (n=2)<br>Wistar (n=1) | Male (n=3) | 1 (n=3) | 2–7 days (n=2)<br>15–28 days (2–4 weeks) (n=1) |
| HomeCageScan (Cleversys, USA) | 10 | Mouse (n=8)<br>Rat ( n=2) | C57BL/6 (n=1)<br>CBA (n=1)<br>DBA/2 (n=1)<br>NOD (n=1)<br>Swiss Webster (n=1)<br>129 (n=1)<br>C57BL/6 (n=3)<br>Mixed background n=3)<br>Long Evans (n=1)<br>Sprague Dawley (n=2) | Male (n =5)<br>Female (n=1)<br>Both (n=2)<br>N/A (n=2) | 1 (n=10) | 2–7 days (n=3)<br>8–14 days (1–2 weeks) (n=1)<br>15–28 days (2–4 weeks) (n=1)<br>29–84 days (1–3 months) (n =4)<br>N/A (n=1) |
| iButtons (Maxim Integrated Products, United Kingdom) | 1 | Mouse (n=1) | C57BL/6 (n=1)<br>129 (n=1)<br>Mixed background (n=1) | Male (n=1) | 1 (n=1) | 29–84 days (1–3 months) (n =1) |
| InfraMot (TSE, USA/Germany) | 6 | Mouse (n=4)<br>Rat (n=2) | C57BL/6 (n=4)<br>Sprague Dawley (n=2)<br>Wistar (n=1) | Male (n=1)<br>Female (n=1)<br>Both (n=3)<br>N/A (n=1) | 1 (n=5)<br>5 (n=1) | 1 day (n=1)<br>2–7 days (n=2)<br>8–14 days (1–2 weeks) (n=1)<br>15–28 days (2–4 weeks) (n=1)<br>29–84 days (1–3 months) (n =1) |
| Infrared Motion Detector (Starr Life Sciences, USA) | 2 | Rat (n=2) | Sprague Dawley (n=2) | Male (n=1)<br>Both (n=1) | 1 (n=1)<br>6 (n=1) | 8–14 days (1–2 weeks) (n=1)<br>85–168 days (3–6 months) (n=1) |

### S4 Supporting Information – Home cage monitoring system

| | | | | | | |
|---|---|---|---|---|---|---|
| Infrared motion sensors (ClockLab, Actimetrics, USA) | 1 | Mouse (n=1) | Mixed background (n=1) | Male (n=1) | 1 ( n=1) | 29–84 days (1–3 months) (n =1) |
| Intellicage (TSE, USA/Germany) | 9 | Mouse (n=8)<br>Rat (n=1) | C57BL/6 (n=5)<br>BALB/c (n=3)<br>DBA/2 (n=3)<br>129 (n=1)<br>Mixed background (n=3)<br>Sprague Dawley (n=1) | Male (n=2)<br>Female (n=4)<br>Both (n=3) | 4 (n=1)<br>7 (n=1)<br>8 (n =2)<br>10 (n=3)<br>> 10 (n=2) | 2–7 days (n=1)<br>8–14 days (1–2 weeks) (n=2)<br>15–28 days (2–4 weeks) (n=3)<br>29–84 days (1–3 months) (n =1)<br>169–336 days (6–12 months) (n=1)<br>N/A (n=1) |
| LabMaster, Environmental chamber, WB 2000 KHL (TSE, USA/Germany) | 1 | Mouse (n=1) | C57BL/6 (n=1) | Male (n=1) | N/A (n=1) | 15–28 days (2–4 weeks) (n=1) |
| Laboras (Metris, the Netherlands) | 4 | Mouse (n=3)<br>Rat (n=1) | C57BL/6 (n=3)<br>Sprague Dawley (n=1) | Male (n=3)<br>Female (n=1) | 1 (n=3)<br>2 (n=1) | 15–28 days (2–4 weeks) (n=1)<br>29–84 days (1–3 months) (n =2)<br>N/A (n=1) |
| Marlau cage (ViewPoint Behavior Technology) | 1 | Rat (n=1) | Sprague Dawley (n=1) | Male (n=1) | > 10 (n=1) | 85–168 days (3–6 months) (n=1) |
| MicroMax Activity monitor (AccuScan Instruments, USA) | 2 | Mouse (n=2) | C57BL/6 (n=1)<br>CD-1 (n=1) | Male (n=2) | 1 (n=1)<br>4 (n=1) | 1 day (n=1)<br>8–14 days (1–2 weeks) (n=1) |
| Mini Run-a-Round (Pets International, Elk Grove Village, IL, USA) | 1 | Mouse (n=1) | C57BL/6 (n=1)<br>129 (n=1) | Both (n=1) | 1 (n=1) | 85–168 days (3–6 months) (n=1) |
| Nano tag (ACOS, Nagano, Japan) | 1 | Rat (n=1) | Fischer 344 (n=1) | Male (n=1) | 3 (n=1) | 29–84 days (1–3 months) (n =1) |
| Noninvasive sleep-monitoring apparatus (Signal Solutions, Lexington, KY) | 1 | Mouse (n=1) | BALB/c (n=1)<br>C57BL/6 (n=1)<br>CD-1 (n=1) | Both (n=1) | 1 (n=1) | 15–28 days (2–4 weeks) (n=1) |
| Operant chambers (Coulbourn Instruments, USA) | 2 | Rat (n=2) | Sprague Dawley (n=2) | Male (n=2) | 1 (n=2) | 8–14 days (1–2 weeks) (n=1)<br>15–28 days (2–4 weeks) (n=1) |
| Operant chambers (MED Associates, USA) | 3 | Mouse (n=1)<br>Rat (n=2) | C57BL/6 (n=1)<br>DBA/2 (n=1) | Male (n=3) | 1 (n=3) | 8–14 days (1–2 weeks) (n=2)<br>N/A (n=1) |

### S4 Supporting Information – Home cage monitoring system

| | | | | | | |
|---|---|---|---|---|---|---|
| | | | Long Evans (n=1)<br>MR (n=1)<br>MNRA (n=1) | | | |
| Opto-M3 Dual Axis System (Columbus Instruments, USA) | 4 | Mouse (n=1)<br>Rat (n=3) | C57BL/6 (n=1)<br>Lister hooded (n=2)<br>Sprague Dawley (n=1) | Male (n=3)<br>Both (n=1) | 1 (n=4) | 2–7 days (n=2)<br>15–28 days (2–4 weeks) (n=1)<br>N/A (n=1) |
| PhenoMaster (TSE, USA/Germany) | 7 | Mouse (n=5)<br>Rat (n=2) | C57BL/6 (n=4)<br>DBA/2 (n=1)<br>129 (n=1)<br>C3H (n=1)<br>Mixed background (n=2)<br>Long Evans (n=1)<br>Sprague Dawley (n=2) | Male (n=3)<br>Female (n=1)<br>Both (n=3) | 1 (n=6)<br>10 (n=1) | 2–7 days (n=3)<br>15–28 days (2–4 weeks) (n=1)<br>29–84 days (1–3 months) (n =1)<br>85–168 days (3–6 months) (n=1)<br>169–336 days (6–12 months) (n=1) |
| Phenopy | 1 | | | | | |
| Phenotyper (Noldus, the Netherlands) | 26 | Mouse (n=23)<br>Rat (n=3) | C57BL/6 (n=21)<br>DBA/2 (n=8)<br>129 (n=4)<br>A/J (n=3)<br>BTBR (n=2)<br>FVB (n=2)<br>AKR (n=1)<br>BALB/c (n=2)<br>C3H (n=2)<br>NOD (n=2)<br>WSB (n=1)<br>PWK (n=1)<br>CAST (n=1)<br>Mixed background (n=2)<br>Sprague Dawley (n=2)<br>Wistar (n=1) | Male (n=15)<br>Female (n=2)<br>Both (n=8)<br>N/A (n=1) | 1 (n=16)<br>2 (n=2)<br>3 (n=1)<br>4 (n=3)<br>5 (n=1)<br>10 (n=1)<br>N/A (n=2) | 2–7 days (n=13)<br>8–14 days (1–2 weeks) (n=4)<br>15–28 days (2–4 weeks) (n=6)<br>29–84 days (1–3 months) (n =3) |
| PhenoWorld (TSE, USA/Germany) | 1 | Rat (n=1) | Wistar (n=1) | Male (n=1) | 6 (n=1) | 29–84 days (1–3 months) (n =1) |
| Photobeam Activity System (Columbus Instruments, USA) | 1 | Mouse (n=1) | C57BL/6 (n=1) | Male (n=1) | 1 (n=1) | N/A (n=1) |

#### S4 Supporting Information – Home cage monitoring system

| | | | | | | |
|---|---|---|---|---|---|---|
| Photobeam Activity System-Home Cage (San Diego Instruments) | 3 | Mouse (n=3) | C57BL/6 (n=2)<br>Mixed background (n=1)<br>129 (n=1) | Both (n=1)<br>Male (n=2) | 1 (n=3) | N/A (n=1)<br>8–14 days (1–2 weeks) (n=1)<br>29–84 days (1–3 months) (n =1) |
| Physiobelt system (Open Science, Russia) | 1 | Rat (n=1) | Long Evans (n=1) | Male (n=1) | 1 (n=1) | 2–7 days (n=1) |
| Radiotelemeter catheter (DSI, USA) | 1 | Mouse (n=1) | C57BL/6 (n=1) | Male (n=1( | N/A (n=1) | 2–7 days (n=1) |
| SAMAB (System for Automated Measurement of Behaviour) | 1 | Mouse (n=1) | C57BL/6 (n=1) | Male (n=1) | 1 (n=1) | 29–84 days (1–3 months) (n =1) |
| Selective Activity Meter (Model S) (Columbus Instruments, USA) | 1 | Mouse (n=1) | OF-1 (n=1) | Male (n=1) | 6 (n=1) | 15–28 days (2–4 weeks) (n=1) |
| SmartFrame (Lafayette Instrument, USA) | 1 | Mouse (n=1) | FVB (n=1) | Male (n=1) | 1 (n=1) | 29–84 days (1–3 months) (n =1) |
| TopScan (CleverSys, USA) | 1 | Mouse (n=1) | C57BL/6 (n=1) | Male (n=1) | 1 (n=1) | 8–14 days (1–2 weeks) (n=1) |
| Videotrack (O'Hara & Co., Tokyo, Japan) | 1 | Mouse (n=1) | C57BL/6 (n=1) | Male (n=1) | 2 (n=1) | 2–7 days (n=1) |
| Vium Digital Smart Houses (San Mateo, CA , USA) | 1 | Rat (n=1) | Lewis (n=1) | Male (n=1) | 2 (n=1) | 15–28 days (2–4 weeks) (n=1) |
| Winchester (5 led), EEG recording system (2 Grass, 12 tracks, 79D model), rotating collector (APCL 12 channels, Air precision) | 1 | Mouse (n=1) | Swiss Webster (n=1) | Male (n=1) | 1 (n=1) | 15–28 days (2–4 weeks) (n=1) |
