## Supplementary material for "Development and Application of Home Cage Monitoring in Laboratory Mice and Rats: a Systematic Review": S5 Supporting information

### S5 Supporting Information – Behavioral, physiological, and external appearance-related parameters

| Degree of automatization | | Monitoring of individuals or groups | | Duration of measurement per day | | Overall duration of measurement | |
|---|---|---|---|---|---|---|---|
| **BEHAVIORAL PARAMETERS** | | | | | | | |
| **Locomotor activity** | | | | | | | |
| Automatic | 222 | Individuals* | 279 | 24 hours | 175 | 1 day | 54 |
| Manual | 89 | Groups* | 32 | < 24 hours, > 12 hours | 18 | 2–7 days | 119 |
| N/A | 4 | N/A | 5 | < 12 hours | 115 | 8–14 days (1–2 weeks) | 39 |
| (*In one study, both individuals and groups were monitored) | | | | N/A | 7 | 15–28 days (2–4 weeks) | 31 |
| | | | | | | 29–84 days (1–3 months) | 37 |
| | | | | | | 85–168 days (3–6 months) | 8 |
| | | | | | | 169–336 days (6–12 months) | 0 |
| | | | | | | 337 and more days (> 1 year) | 1 |
| | | | | | | N/A | 26 |
| **Feeding (drinking, food)** | | | | | | | |
| Automatic | 78 | Individuals | 178 | 24 hours | 115 | 1 day | 20 |
| Manual | 127 | Groups | 28 | < 24 hours, > 12 hours | 13 | 2–7 days | 57 |
| N/A | 6 | N/A | 5 | < 12 hours | 74 | 8–14 days (1–2 weeks) | 35 |
| | | | | N/A | 9 | 15–28 days (2–4 weeks) | 37 |
| | | | | | | 29–84 days (1–3 months) | 35 |
| | | | | | | 85–168 days (3–6 months) | 6 |
| | | | | | | 169–336 days (6–12 months) | 1 |
| | | | | | | 337 and more days (> 1 year) | 1 |
| | | | | | | N/A | 19 |
| **Social behavior** | | | | | | | |
| Automatic | 14 | Individuals | 83 | 24 hours | 14 | 1 day | 27 |
| Manual | 101 | Groups | 34 | < 24 hours, > 12 hours | 2 | 2–7 days | 39 |
| N/A | 2 | N/A | | < 12 hours | 100 | 8–14 days (1–2 weeks) | 19 |
| | | | | N/A | 1 | 15–28 days (2–4 weeks) | 11 |
| | | | | | | 29–84 days (1–3 months) | 10 |
| | | | | | | 85–168 days (3–6 months) | 4 |
| | | | | | | 169–336 days (6–12 months) | 1 |
| | | | | | | 337 and more days (> 1 year) | 1 |
| | | | | | | N/A | 5 |
| **Burrowing and nesting** | | | | | | | |
| Automatic | 8 | Individuals | 44 | 24 hours | 10 | 1 day | 17 |
| Manual | 46 | Groups | 10 | < 24 hours, > 12 hours | 3 | 2–7 days | 19 |
| N/A | 2 | N/A | 2 | < 12 hours | 43 | 8–14 days (1–2 weeks) | 11 |
| | | | | N/A | | 15–28 days (2–4 weeks) | 4 |
| | | | | | | 29–84 days (1–3 months) | 3 |
| | | | | | | 85–168 days (3–6 months) | |
| | | | | | | 169–336 days (6–12 months) | |
| | | | | | | 337 and more days (> 1 year) | 1 |
| | | | | | | N/A | 1 |

### S5 Supporting Information – Behavioral, physiological, and external appearance-related parameters

| | | | | | | | |
|---|---|---|---|---|---|---|---|
| **Wheel running** | | | | | | | |
| Automatic | 47 | Individuals | 45 | 24 hours | 44 | 1 day | 2 |
| Manual | 2 | Groups | 2 | < 24 hours, > 12 hours | 2 | 2–7 days | 11 |
| N/A | | N/A | 2 | < 12 hours | 1 | 8–14 days (1–2 weeks) | 8 |
| | | | | N/A | 2 | 15–28 days (2–4 weeks) | 11 |
| | | | | | | 29–84 days (1–3 months) | 12 |
| | | | | | | 85–168 days (3–6 months) | 2 |
| | | | | | | 169–336 days (6–12 months) | 1 |
| | | | | | | 337 and more days (> 1 year) | |
| | | | | | | N/A | 2 |
| **Abnormal behaviors** | | | | | | | |
| Automatic | 3 | Individuals | 27 | 24 hours | 5 | 1 day | 3 |
| Manual | 29 | Groups | 6 | < 24 hours, > 12 hours | 3 | 2–7 days | 12 |
| N/A | 1 | N/A | | < 12 hours | 24 | 8–14 days (1–2 weeks) | 4 |
| | | | | N/A | 1 | 15–28 days (2–4 weeks) | 1 |
| | | | | | | 29–84 days (1–3 months) | 6 |
| | | | | | | 85–168 days (3–6 months) | 1 |
| | | | | | | 169–336 days (6–12 months) | 3 |
| | | | | | | 337 and more days (> 1 year) | |
| | | | | | | N/A | 3 |
| **Facial expression and body posture** | | | | | | | |
| Automatic | 4 | Individuals | 25 | 24 hours | 5 | 1 day | 5 |
| Manual | 24 | Groups | 2 | < 24 hours, > 12 hours | 1 | 2–7 days | 9 |
| N/A | 1 | N/A | 2 | < 12 hours | 21 | 8–14 days (1–2 weeks) | 3 |
| | | | | N/A | 2 | 15–28 days (2–4 weeks) | 4 |
| | | | | | | 29–84 days (1–3 months) | 5 |
| | | | | | | 85–168 days (3–6 months) | 1 |
| | | | | | | 169–336 days (6–12 months) | |
| | | | | | | 337 and more days (> 1 year) | |
| | | | | | | N/A | 2 |
| **Grooming** | | | | | | | |
| Automatic | 7 | Individuals | 24 | 24 hours | 3 | 1 day | 9 |
| Manual | 19 | Groups | 2 | < 24 hours, > 12 hours | 1 | 2–7 days | 8 |
| N/A | 1 | N/A | 1 | < 12 hours | 21 | 8–14 days (1–2 weeks) | 3 |
| | | | | N/A | 2 | 15–28 days (2–4 weeks) | 1 |
| | | | | | | 29–84 days (1–3 months) | 2 |
| | | | | | | 85–168 days (3–6 months) | |
| | | | | | | 169–336 days (6–12 months) | |
| | | | | | | 337 and more days (> 1 year) | |
| | | | | | | N/A | 4 |

### S5 Supporting Information – Behavioral, physiological, and external appearance-related parameters

| | | | | | | | |
|---|---|---|---|---|---|---|---|
| **Learning and memory** | | | | | | | |
| Automatic | 16 | Individuals | 23 | 24 hours | 12 | 1 day | 1 |
| Manual | 6 | Groups | 1 | < 24 hours, > 12 hours | 1 | 2–7 days | 7 |
| N/A | 2 | N/A | | < 12 hours | 10 | 8–14 days (1–2 weeks) | 8 |
| | | | | N/A | 1 | 15–28 days (2–4 weeks) | 3 |
| | | | | | | 29–84 days (1–3 months) | 4 |
| | | | | | | 85–168 days (3–6 months) | 1 |
| | | | | | | 169–336 days (6–12 months) | |
| | | | | | | 337 and more days (> 1 year) | |
| | | | | | | N/A | |
| **Anxiety, depression, and schizophrenia** | | | | | | | |
| Automatic | 4 | Individuals | 14 | 24 hours | 4 | 1 day | 2 |
| Manual | 10 | Groups | | < 24 hours, > 12 hours | | 2–7 days | 6 |
| N/A | | N/A | | < 12 hours | 9 | 8–14 days (1–2 weeks) | 2 |
| | | | | N/A | 1 | 15–28 days (2–4 weeks) | 1 |
| | | | | | | 29–84 days (1–3 months) | 2 |
| | | | | | | 85–168 days (3–6 months) | |
| | | | | | | 169–336 days (6–12 months) | |
| | | | | | | 337 and more days (> 1 year) | |
| | | | | | | N/A | 1 |
| **Sleep behavior** | | | | | | | |
| Automatic | 6 | Individuals | 11 | 24 hours | 6 | 1 day | 4 |
| Manual | 6 | Groups | 1 | < 24 hours, > 12 hours | 1 | 2–7 days | 3 |
| N/A | | N/A | | < 12 hours | 5 | 8–14 days (1–2 weeks) | 1 |
| | | | | N/A | | 15–28 days (2–4 weeks) | 3 |
| | | | | | | 29–84 days (1–3 months) | |
| | | | | | | 85–168 days (3–6 months) | |
| | | | | | | 169–336 days (6–12 months) | |
| | | | | | | 337 and more days (> 1 year) | |
| | | | | | | N/A | 1 |
| **Vocalisation** | | | | | | | |
| Automatic | 3 | Individuals | 8 | 24 hours | | 1 day | 4 |
| Manual | 6 | Groups | 2 | < 24 hours, > 12 hours | 1 | 2–7 days | 6 |
| N/A | 2 | N/A | 1 | < 12 hours | 10 | 8–14 days (1–2 weeks) | 1 |
| | | | | N/A | | 15–28 days (2–4 weeks) | |
| | | | | | | 29–84 days (1–3 months) | |
| | | | | | | 85–168 days (3–6 months) | |
| | | | | | | 169–336 days (6–12 months) | |
| | | | | | | 337 and more days (> 1 year) | |
| | | | | | | N/A | |

### S5 Supporting Information – Behavioral, physiological, and external appearance-related parameters

| | | | | | | | |
|---|---|---|---|---|---|---|---|
| **Motor and sensory functions** | | | | | | | |
| Automatic | 6 | Individuals | 8 | 24 hours | 4 | 1 day | |
| Manual | 3 | Groups | 1 | < 24 hours, > 12 hours | 1 | 2–7 days | 1 |
| N/A | | N/A | | < 12 hours | 3 | 8–14 days (1–2 weeks) | 1 |
| | | | | N/A | 1 | 15–28 days (2–4 weeks) | 3 |
| | | | | | | 29–84 days (1–3 months) | 1 |
| | | | | | | 85–168 days (3–6 months) | |
| | | | | | | 169–336 days (6–12 months) | |
| | | | | | | 337 and more days (> 1 year) | |
| | | | | | | N/A | 3 |
| **Spatial preference** | | | | | | | |
| Automatic | 4 | Individuals | 4 | 24 hours | 4 | 1 day | |
| Manual | 2 | Groups | 2 | < 24 hours, > 12 hours | 1 | 2–7 days | 4 |
| N/A | | N/A | | < 12 hours | 1 | 8–14 days (1–2 weeks) | 2 |
| | | | | N/A | | 15–28 days (2–4 weeks) | |
| | | | | | | 29–84 days (1–3 months) | |
| | | | | | | 85–168 days (3–6 months) | |
| | | | | | | 169–336 days (6–12 months) | |
| | | | | | | 337 and more days (> 1 year) | |
| | | | | | | N/A | |
| **Defecation and urination** | | | | | | | |
| Automatic | | Individuals | 4 | 24 hours | 3 | 1 day | 1 |
| Manual | 5 | Groups | 1 | < 24 hours, > 12 hours | | 2–7 days | 1 |
| N/A | | N/A | | < 12 hours | 2 | 8–14 days (1–2 weeks) | 1 |
| | | | | N/A | | 15–28 days (2–4 weeks) | |
| | | | | | | 29–84 days (1–3 months) | 1 |
| | | | | | | 85–168 days (3–6 months) | |
| | | | | | | 169–336 days (6–12 months) | |
| | | | | | | 337 and more days (> 1 year) | |
| | | | | | | N/A | 1 |
| **Sniffing** | | | | | | | |
| Automatic | 1 | Individuals | 4 | 24 hours | | 1 day | 2 |
| Manual | 2 | Groups | | < 24 hours, > 12 hours | | 2–7 days | |
| N/A | 1 | N/A | | < 12 hours | 3 | 8–14 days (1–2 weeks) | |
| | | | | N/A | 1 | 15–28 days (2–4 weeks) | 1 |
| | | | | | | 29–84 days (1–3 months) | |
| | | | | | | 85–168 days (3–6 months) | |
| | | | | | | 169–336 days (6–12 months) | |
| | | | | | | 337 and more days (> 1 year) | |
| | | | | | | N/A | 1 |

### S5 Supporting Information – Behavioral, physiological, and external appearance-related parameters

| | | | | | | | |
|---|---|---|---|---|---|---|---|
| **Seizures** | | | | | | | |
| Automatic | 1 | Individuals | 3 | 24 hours | 1 | 1 day | |
| Manual | 2 | Groups | | < 24 hours, > 12 hours | 1 | 2–7 days | 2 |
| N/A | | N/A | | < 12 hours | 1 | 8–14 days (1–2 weeks) | 1 |
| | | | | N/A | | 15–28 days (2–4 weeks) | |
| | | | | | | 29–84 days (1–3 months) | |
| | | | | | | 85–168 days (3–6 months) | |
| | | | | | | 169–336 days (6–12 months) | |
| | | | | | | 337 and more days (> 1 year) | |
| | | | | | | N/A | |
| Automatic | 1 | Individuals | 3 | 24 hours | 1 | 1 day | |
| **Champing behavior** | | | | | | | |
| Automatic | | Individuals | | 24 hours | 1 | 1 day | |
| Manual | 1 | Groups | 1 | < 24 hours, > 12 hours | | 2–7 days | |
| N/A | | N/A | | < 12 hours | | 8–14 days (1–2 weeks) | |
| | | | | N/A | | 15–28 days (2–4 weeks) | |
| | | | | | | 29–84 days (1–3 months) | |
| | | | | | | 85–168 days (3–6 months) | |
| | | | | | | 169–336 days (6–12 months) | |
| | | | | | | 337 and more days (> 1 year) | 1 |
| | | | | | | N/A | |
| **Clinical sings not further specified** | | | | | | | |
| Automatic | | Individuals | 1 | 24 hours | | 1 day | |
| Manual | 1 | Groups | | < 24 hours, > 12 hours | | 2–7 days | |
| N/A | | N/A | | < 12 hours | | 8–14 days (1–2 weeks) | |
| | | | | N/A | 1 | 15–28 days (2–4 weeks) | 1 |
| | | | | | | 29–84 days (1–3 months) | |
| | | | | | | 85–168 days (3–6 months) | |
| | | | | | | 169–336 days (6–12 months) | |
| | | | | | | 337 and more days (> 1 year) | |
| | | | | | | N/A | |
| **Colonic contractility** | | | | | | | |
| Automatic | 1 | Individuals | 1 | 24 hours | | 1 day | |
| Manual | | Groups | | < 24 hours, > 12 hours | | 2–7 days | 1 |
| N/A | | N/A | | < 12 hours | 1 | 8–14 days (1–2 weeks) | |
| | | | | N/A | | 15–28 days (2–4 weeks) | |
| | | | | | | 29–84 days (1–3 months) | |
| | | | | | | 85–168 days (3–6 months) | |
| | | | | | | 169–336 days (6–12 months) | |
| | | | | | | 337 and more days (> 1 year) | |
| | | | | | | N/A | |

### S5 Supporting Information – Behavioral, physiological, and external appearance-related parameters

| | | | | | | | |
|---|---|---|---|---|---|---|---|
| **Curiosity/altertness** | | | | | | | |
| Automatic | | Individuals | 1 | 24 hours | | 1 day | |
| Manual | 1 | Groups | | < 24 hours, > 12 hours | | 2–7 days | 1 |
| N/A | | N/A | | < 12 hours | | 8–14 days (1–2 weeks) | |
| | | | | N/A | 1 | 15–28 days (2–4 weeks) | |
| | | | | | | 29–84 days (1–3 months) | |
| | | | | | | 85–168 days (3–6 months) | |
| | | | | | | 169–336 days (6–12 months) | |
| | | | | | | 337 and more days (> 1 year) | |
| | | | | | | N/A | |
| **Sign of pain or distress not further specified** | | | | | | | |
| Automatic | | Individuals | 1 | 24 hours | | 1 day | |
| Manual | 1 | Groups | | < 24 hours, > 12 hours | | 2–7 days | 1 |
| N/A | | N/A | | < 12 hours | | 8–14 days (1–2 weeks) | |
| | | | | N/A | 1 | 15–28 days (2–4 weeks) | |
| | | | | | | 29–84 days (1–3 months) | |
| | | | | | | 85–168 days (3–6 months) | |
| | | | | | | 169–336 days (6–12 months) | |
| | | | | | | 337 and more days (> 1 year) | |
| | | | | | | N/A | |
| **Twitches** | | | | | | | |
| Automatic | 1 | Individuals | 1 | 24 hours | 1 | 1 day | |
| Manual | | Groups | | < 24 hours, > 12 hours | | 2–7 days | 1 |
| N/A | | N/A | | < 12 hours | | 8–14 days (1–2 weeks) | |
| | | | | N/A | | 15–28 days (2–4 weeks) | |
| | | | | | | 29–84 days (1–3 months) | |
| | | | | | | 85–168 days (3–6 months) | |
| | | | | | | 169–336 days (6–12 months) | |
| | | | | | | 337 and more days (> 1 year) | |
| | | | | | | N/A | |
| **Other: behavior not further specified** | | | | | | | |
| Automatic | 1 | Individuals | | 24 hours | | 1 day | |
| Manual | | Groups | | < 24 hours, > 12 hours | | 2–7 days | 1 |
| N/A | | N/A | 1 | < 12 hours | 1 | 8–14 days (1–2 weeks) | |
| | | | | N/A | | 15–28 days (2–4 weeks) | |
| | | | | | | 29–84 days (1–3 months) | |
| | | | | | | 85–168 days (3–6 months) | |
| | | | | | | 169–336 days (6–12 months) | |
| | | | | | | 337 and more days (> 1 year) | |
| | | | | | | N/A | |

### S5 Supporting Information – Behavioral, physiological, and external appearance-related parameters

| | | | | | | | |
|---|---|---|---|---|---|---|---|
| **PHYSIOLOGICAL PARAMETERS** | | | | | | | |
| **Body temperature** | | | | | | | |
| Automatic | 34 | Individuals | 36 | 24 hours | 25 | 1 day | 9 |
| Manual | 3 | Groups | 1 | < 24 hours, > 12 hours | 2 | 2–7 days | 11 |
| N/A | | N/A | | < 12 hours | 9 | 8–14 days (1–2 weeks) | 6 |
| | | | | N/A | 1 | 15–28 days (2–4 weeks) | 3 |
| | | | | | | 29–84 days (1–3 months) | 5 |
| | | | | | | 85–168 days (3–6 months) | |
| | | | | | | 169–336 days (6–12 months) | |
| | | | | | | 337 and more days (> 1 year) | 1 |
| | | | | | | N/A | 2 |
| **Body weight** | | | | | | | |
| Automatic | 2 | Individuals | 2 | 24 hours | 2 | 1 day | |
| Manual | | Groups | | < 24 hours, > 12 hours | | 2–7 days | 1 |
| N/A | | N/A | | < 12 hours | | 8–14 days (1–2 weeks) | |
| | | | | N/A | | 15–28 days (2–4 weeks) | |
| | | | | | | 29–84 days (1–3 months) | 1 |
| | | | | | | 85–168 days (3–6 months) | |
| | | | | | | 169–336 days (6–12 months) | |
| | | | | | | 337 and more days (> 1 year) | |
| | | | | | | N/A | |
| **Electroencephalography** | | | | | | | |
| Automatic | 8 | Individuals | 8 | 24 hours | 6 | 1 day | 5 |
| Manual | | Groups | | < 24 hours, > 12 hours | 1 | 2–7 days | 2 |
| N/A | | N/A | | < 12 hours | 1 | 8–14 days (1–2 weeks) | 1 |
| | | | | N/A | | 15–28 days (2–4 weeks) | |
| | | | | | | 29–84 days (1–3 months) | |
| | | | | | | 85–168 days (3–6 months) | |
| | | | | | | 169–336 days (6–12 months) | |
| | | | | | | 337 and more days (> 1 year) | |
| | | | | | | N/A | |
| **Electromyography** | | | | | | | |
| Automatic | 7 | Individuals | 7 | 24 hours | 5 | 1 day | 5 |
| Manual | | Groups | | < 24 hours, > 12 hours | | 2–7 days | 1 |
| N/A | | N/A | | < 12 hours | 2 | 8–14 days (1–2 weeks) | |
| | | | | N/A | | 15–28 days (2–4 weeks) | |
| | | | | | | 29–84 days (1–3 months) | |
| | | | | | | 85–168 days (3–6 months) | |
| | | | | | | 169–336 days (6–12 months) | |
| | | | | | | 337 and more days (> 1 year) | |
| | | | | | | N/A | 1 |

### S5 Supporting Information – Behavioral, physiological, and external appearance-related parameters

| | | | | | | | |
|---|---|---|---|---|---|---|---|
| **Heart rate & Electrocardiography** | | | | | | | |
| Automatic | 46 | Individuals | 49 | 24 hours | 24 | 1 day | 8 |
| Manual | 3 | Groups | | < 24 hours, > 12 hours | 4 | 2–7 days | 14 |
| N/A | | N/A | | < 12 hours | 20 | 8–14 days (1–2 weeks) | 9 |
| | | | | N/A | 1 | 15–28 days (2–4 weeks) | 6 |
| | | | | | | 29–84 days (1–3 months) | 5 |
| | | | | | | 85–168 days (3–6 months) | |
| | | | | | | 169–336 days (6–12 months) | |
| | | | | | | 337 and more days (> 1 year) | |
| | | | | | | N/A | 7 |
| **Blood pressure** | | | | | | | |
| Automatic | 25 | Individuals | 26 | 24 hours | 13 | 1 day | 3 |
| Manual | 1 | Groups | | < 24 hours, > 12 hours | 3 | 2–7 days | 5 |
| N/A | | N/A | | < 12 hours | 10 | 8–14 days (1–2 weeks) | 5 |
| | | | | N/A | | 15–28 days (2–4 weeks) | 5 |
| | | | | | | 29–84 days (1–3 months) | 4 |
| | | | | | | 85–168 days (3–6 months) | |
| | | | | | | 169–336 days (6–12 months) | |
| | | | | | | 337 and more days (> 1 year) | |
| | | | | | | N/A | 4 |
| **Respiration** | | | | | | | |
| Automatic | 4 | Individuals | 8 | 24 hours | 3 | 1 day | 1 |
| Manual | 4 | Groups | | < 24 hours, > 12 hours | | 2–7 days | 4 |
| N/A | | N/A | | < 12 hours | 5 | 8–14 days (1–2 weeks) | 3 |
| | | | | N/A | | 15–28 days (2–4 weeks) | |
| | | | | | | 29–84 days (1–3 months) | |
| | | | | | | 85–168 days (3–6 months) | |
| | | | | | | 169–336 days (6–12 months) | |
| | | | | | | 337 and more days (> 1 year) | |
| | | | | | | N/A | |
| **(Stress) hormones** | | | | | | | |
| Automatic | 2 | Individuals | 5 | 24 hours | 1 | 1 day | 2 |
| Manual | 4 | Groups | 2 | < 24 hours, > 12 hours | 1 | 2–7 days | 3 |
| N/A | 1 | N/A | | < 12 hours | 5 | 8–14 days (1–2 weeks) | |
| | | | | N/A | | 15–28 days (2–4 weeks) | 1 |
| | | | | | | 29–84 days (1–3 months) | 1 |
| | | | | | | 85–168 days (3–6 months) | |
| | | | | | | 169–336 days (6–12 months) | |
| | | | | | | 337 and more days (> 1 year) | |
| | | | | | | N/A | |

### S5 Supporting Information – Behavioral, physiological, and external appearance-related parameters

| | | | | | | | |
|---|---|---|---|---|---|---|---|
| **Neuronal activity** | | | | | | | |
| Automatic | 12 | Individuals | 13 | 24 hours | 7 | 1 day | 2 |
| Manual | 1 | Groups | | < 24 hours, > 12 hours | | 2–7 days | 5 |
| N/A | | N/A | | < 12 hours | 5 | 8–14 days (1–2 weeks) | 1 |
| | | | | N/A | 1 | 15–28 days (2–4 weeks) | 1 |
| | | | | | | 29–84 days (1–3 months) | 1 |
| | | | | | | 85–168 days (3–6 months) | 1 |
| | | | | | | 169–336 days (6–12 months) | |
| | | | | | | 337 and more days (> 1 year) | |
| | | | | | | N/A | 2 |
| **EXTERNAL APPEARANCE** | | | | | | | |
| **Body Condition Score** | | | | | | | |
| Automatic | | Individuals | 1 | 24 hours | | 1 day | |
| Manual | 4 | Groups | 2 | < 24 hours, > 12 hours | | 2–7 days | 1 |
| N/A | | N/A | 1 | < 12 hours | 2 | 8–14 days (1–2 weeks) | |
| | | | | N/A | | 15–28 days (2–4 weeks) | 1 |
| | | | | | | 29–84 days (1–3 months) | |
| | | | | | | 85–168 days (3–6 months) | 1 |
| | | | | | | 169–336 days (6–12 months) | 1 |
| | | | | | | 337 and more days (> 1 year) | |
| | | | | | | N/A | |
| **Fur condition** | | | | | | | |
| Automatic | | Individuals | 3 | 24 hours | 1 | 1 day | |
| Manual | 6 | Groups | 2 | < 24 hours, > 12 hours | | 2–7 days | 2 |
| N/A | | N/A | 1 | < 12 hours | 3 | 8–14 days (1–2 weeks) | |
| | | | | N/A | 2 | 15–28 days (2–4 weeks) | 1 |
| | | | | | | 29–84 days (1–3 months) | 1 |
| | | | | | | 85–168 days (3–6 months) | |
| | | | | | | 169–336 days (6–12 months) | 2 |
| | | | | | | 337 and more days (> 1 year) | |
| | | | | | | N/A | |
| **Piloerection** | | | | | | | |
| Automatic | | Individuals | 3 | 24 hours | | 1 day | |
| Manual | 3 | Groups | | < 24 hours, > 12 hours | | 2–7 days | 2 |
| N/A | 1 | N/A | 1 | < 12 hours | 3 | 8–14 days (1–2 weeks) | |
| | | | | N/A | 1 | 15–28 days (2–4 weeks) | |
| | | | | | | 29–84 days (1–3 months) | 1 |
| | | | | | | 85–168 days (3–6 months) | |
| | | | | | | 169–336 days (6–12 months) | |
| | | | | | | 337 and more days (> 1 year) | |
| | | | | | | N/A | 1 |

### S5 Supporting Information – Behavioral, physiological, and external appearance-related parameters

| | | | | | | | |
|---|---|---|---|---|---|---|---|
| **Wounds** | | | | | | | |
| Automatic | | Individuals | 2 | 24 hours | 1 | 1 day | |
| Manual | 6 | Groups | 4 | < 24 hours, > 12 hours | | 2–7 days | 1 |
| N/A | | N/A | | < 12 hours | 3 | 8–14 days (1–2 weeks) | |
| | | | | N/A | 2 | 15–28 days (2–4 weeks) | |
| | | | | | | 29–84 days (1–3 months) | |
| | | | | | | 85–168 days (3–6 months) | 1 |
| | | | | | | 169–336 days (6–12 months) | 1 |
| | | | | | | 337 and more days (> 1 year) | 2 |
| | | | | | | N/A | 1 |
| **Chromodacryorrhea** | | | | | | | |
| Automatic | | Individuals | 1 | 24 hours | | 1 day | |
| Manual | 1 | Groups | | < 24 hours, > 12 hours | | 2–7 days | |
| N/A | | N/A | | < 12 hours | 1 | 8–14 days (1–2 weeks) | |
| | | | | N/A | | 15–28 days (2–4 weeks) | |
| | | | | | | 29–84 days (1–3 months) | |
| | | | | | | 85–168 days (3–6 months) | 1 |
| | | | | | | 169–336 days (6–12 months) | |
| | | | | | | 337 and more days (> 1 year) | |
| | | | | | | N/A | |
| **External appearance related parameters not further specified** | | | | | | | |
| Automatic | 1 | Individuals | 3 | 24 hours | 2 | 1 day | |
| Manual | 2 | Groups | | < 24 hours, > 12 hours | | 2–7 days | |
| N/A | | N/A | | < 12 hours | | 8–14 days (1–2 weeks) | |
| | | | | N/A | 1 | 15–28 days (2–4 weeks) | |
| | | | | | | 29–84 days (1–3 months) | 2 |
| | | | | | | 85–168 days (3–6 months) | |
| | | | | | | 169–336 days (6–12 months) | |
| | | | | | | 337 and more days (> 1 year) | |
| | | | | | | N/A | 1 |
